# Context-dependent functional compensation between Ythdf m6A readers

**DOI:** 10.1101/2020.06.03.131441

**Authors:** Lior Lasman, Vladislav Krupalnik, Shay Geula, Mirie Zerbib, Sergey Viukov, Nofar Mor, Alejandro Aguilera Castrejon, Orel Mizrahi, Sathe Shashank, Aharon Nachshon, Dan Schneir, Stefan Aigner, Archana Shankar, Jasmine Mueller, Noam Stern-Ginossar, Gene W Yeo, Noa Novershtern, Jacob H Hanna

## Abstract

The N6-methyladenosine (m^6^A) modification is the most prevalent post-transcriptional mRNA modification, regulating mRNA decay, translation and splicing. It plays a major role during normal development, differentiation, and disease progression. The modification is dynamically regulated by a set of writer, eraser and reader proteins. The YTH-domain family of proteins: Ythdf1, Ythdf2, and Ythdf3, are three homologous m^6^A binding proteins, which have different cellular functions. However, their sequence similarity and their tendency to bind the same targets suggest that they may have overlapping roles. We systematically knocked out (KO) the Mettl3 writer for each of the Ythdf readers and for the three readers together (triple-KO). We then estimated the effect *in-vivo*, in mouse gametogenesis and viability, and *in-vitro*, in mouse embryonic stem cells (mESCs). We show that in gametogenesis, Mettl3-KO severity is increased as the deletion occurs earlier in the process, and Ythdf2 has a dominant role that cannot be compensated by Ythdf1 or Ythdf3, possibly due to differences in readers’ expression, both in quantity and in spatial location. By knocking out the three readers together and systematically testing offspring genotypes, we have revealed a redundancy in the readers’ role during early development, a redundancy which is dosage-dependent. Additionally, we show that in mESCs there is compensation between the three readers, since the inability to differentiate and the significant effect on mRNA decay occur only in the triple-KO cells and not in the single KOs. Thus, we suggest a novel model for the Ythdf readers function. There is a dosage-dependent redundancy when all three readers are co-expressed in the same location in the cells.

## Introduction

RNA modifications are a layer of gene expression regulation, similar to DNA and protein modifications (Heck and Wilusz 2019). N6-methyladenosine, also known as m^6^A, is the most abundant mRNA modification (Heck and Wilusz 2019). It was first discovered in the 70’s, but major progress was done in recent years due to new approaches of mapping m^6^A sites (Dominissini et al. 2012; Meyer et al. 2012; Garcia-Campos et al. 2019). Its importance was shown in a wide range of organisms and processes, from yeast meiosis (Schwartz et al. 2013), sex determination in drosophila (Kan et al. 2017), and up to mammalian early development (Geula et al. 2015), neural development (Wang et al. 2018) and hematopoiesis (Lee et al. 2019).

m^6^A modification is highly regulated by writer, reader and eraser proteins. Mettl3 forms a heterodimer with Mettl14 (Liu et al. 2014), and together with the supporting WTAP protein, catalyzes m^6^A with preference to 3’UTR, 5’UTR, long exons, and near stop codons (Heck and Wilusz 2019). FTO and Alkbh5 are eraser enzymes found in vertebrates (Zheng et al. 2013; Jia et al. 2011), and potentially have distinct role and localization in the cell (Zheng et al. 2013; Wei et al. 2018).

Multiple proteins were identified as m^6^A readers, with the YTH domain-containing proteins (Ythdf and Ythdc) stand out among them. Recent studies link these proteins to different functions in RNA metabolism: Ythdf1 and Ythdf3 were shown to promote translation by recruiting translation initiation factors in HeLa cells (Shi et al. 2017; Wang et al. 2015; Li et al. 2017), Ythdf2 was linked to degradation, partially by recruiting the CCR4-NOT deadenylase complex (Du et al. 2016; Wang et al. 2014), and nuclear Ythdc1 was shown to regulate splicing (Kasowitz et al. 2018; Hartmann et al. 1999). Ythdf1, 2 & 3 share high protein sequence similarity (67-70%, **Figure S1a**). In addition, they have co-evolved during evolution. While Drosophila melanogaster has one copy of Ythdf protein (named Ythdf), vertebrates have three functional proteins (**Figure S1b**), probably generated following duplication events (Pervaiz et al. 2019).

It is not fully clear whether each one of the Ythdf readers fulfills a distinct role. Their sequence similarity, and that they are all localized in the cytoplasm (Wang et al. 2015, 2014; Shi et al. 2017) and share many of their targets (Li et al. 2017; Patil et al. 2016, 2018) indicate partial redundancy. However, knockout (KO) of Ythdf2 alone is sufficient to stop proper oocyte maturation (Ivanova et al. 2017), and a single KO of Ythdf1 or of Ythdf2 causes neural defects (Li et al. 2018; Shi et al. 2018), suggesting that in certain systems, Ythdf readers cannot compensate each other. This however could be a result of differences in expression levels in the different tissues. Comprehensive research into the redundancy between the three Ythdfs has not been conducted. In addition, the effect of knocking out the three readers *in-vivo* has not been described so far.

In this study we use conditional single KOs of Mettl3 and Ythdf1, 2 & 3, and show that both Mettl3 and Ythdf2 are essential for proper gametogenesis, and that mice lacking these proteins are either hypo-fertile or sterile. The severity of the phenotype is increased when the Mettl3 deletion is done earlier in the process. We found that the Ythdf readers have different expression patterns during gametogenesis, which might explain the lack of compensation in this process. In addition, we generated an *in vivo* triple-KO and found that it leads to impaired development as early as E7.5, and to embryonic lethality. By using systematic genotyping of viable offspring, we found that in early development there is compensation between the readers, which is dosage-dependent, i.e. Ythdf2-hetrozygouse mice need to have at least one functional copy of another Ythdf reader to escape mortality. Furthermore, we used mESCs to analyze the function of each Ythdf reader separately, and together. We found that only triple-KO mESCs are not able to differentiate properly, and present a prolonged mRNA degradation rate, similar to the effect shown in Mettl3-KO, while no significant effect is seen in the single-KOs. This suggests that just like in early development, in mouse ESCs, a system in which all the readers are expressed in the same cells and compartment, there is a redundancy between Ythdf readers, which enables compensation in the absence of the other.

## Results

### Mettl3 writer plays an essential role in oogenesis and spermatogenesis

We started by systematically testing the three readers in a specific system *in-vivo*, focusing on spermatogenesis and oogenesis. m^6^A writers Mettl3 and Mettl14 and m^6^A erasers FTO and ALKBH5 were found to be essential for proper gametogenesis in mouse. Their KO typically leads to defective maturation of sperm or ova, and hypofertility (Xu et al. 2017; Lin et al. 2017; Zheng et al. 2013; Tang et al. 2017; Lasman, L, Hanna, JH, Novershtern 2020; Kasowitz et al. 2018). As for m^6^A readers, both Ythdc1 and Ythdc2 have an essential role in gametogenesis. Their KO in spermatogenesis or oogenesis leads to a severe hypofertility phenotype (Hsu et al. 2017; Bailey et al. 2017; Wojtas et al. 2017; Jain et al. 2018; Kasowitz et al. 2018). Knocking out Ythdf2 leads to normal ovulation but an inability to downregulate maternal mRNA. Thus, Ythdf2-KO females are sterile (Ivanova et al. 2017). In contrast, Ythdf2-KO males show normal seminiferous tubule histology (Ivanova et al. 2017). Depletion of Ythdf2 in mouse spermatogonia leads to defective cell morphology and decreases cell proliferation (Huang et al. 2020). However, further examination of all Ythdf readers in spermatogenesis is still required.

We assessed the importance of m^6^A in gametogenesis, both in males and females, using conditional KOs of the m^6^A writer Mettl3 (**Figure S2**). Mettl3^flox/flox^ mice were crossed with mice carrying one of the following Cre constructs: ZP3-Cre which is activated during oogenesis, Stra8-Cre and Prm1-Cre, which are activated during spermatogenesis, and Vasa-Cre which is activated in the early stages of both. Thus, we could test the effect of Mettl3-KO systematically in different time points during gametogenesis, in both males and females.

Mettl3^f/f^Vasa-cre+ KO had a major effect on oocyte development. A dissection of Mettl3^f/f^Vasa-cre+ female mice showed abnormal ovary morphology **(Figure 1a-b)**, and the mice were sterile **(Figure 1c)**. Zp3 is expressed during a later stage of the oocyte maturation, prior to the completion of the first meiosis (Gao et al. 2017a). Accordingly, Mettl3^f/f^Zp3-cre+ female mice showed a normal ovary morphology (**Figure 1d**). However, the mice were sterile **(Figure 1e)**. Flushing oocytes from the oviduct revealed an overall significant low number of oocytes (p-value <0.002, **Figure 1f**). All the flushed oocytes of the KO were stuck at the germinal vesicle (GV) stage and did not reach the two-cell stage upon fertilization attempts **(Figure 1g, S3a)**, meaning that they have not completed the first meiosis. Indeed, immunostaining of tubulin in KO and WT oocytes, showed that KO oocytes were stuck in the GV stage, and did not proceed for GV breakdown and completion of the first meiosis **(Figure 1h)**. The transcriptional profile of Mettl3-cKO and control oocytes revealed a major change in transcription **(Figure S3b)**, including aberrant expression of genes related to oocyte development **(Figure S3c-d, Table S1)**.

**Figure 1.**
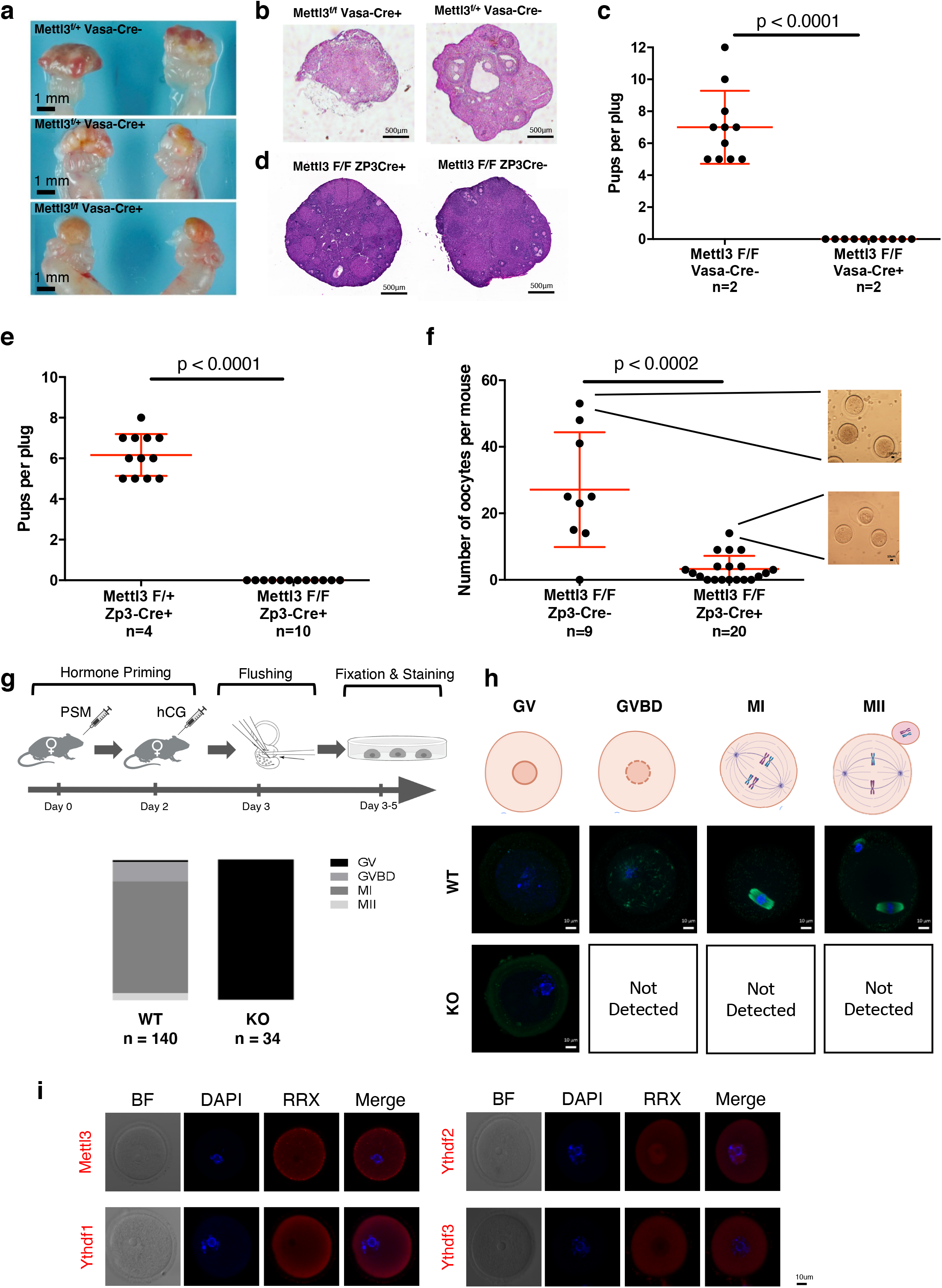
Mettl3 is essential for female mice fertility. **a)** Gross morphology of Cre+ and Cre−(control) female ovaries. Cre+ females show a smooth shape that lacks the typical follicular morphology. **b)** H&E staining of an ovary, showing a severe abnormality in Mettl3^f/f^ Vasa-Cre+ females. **c)** Number of pups per plug produced by mating Mettl3^f/f^ Vasa-Cre+ females, compared to Mettl3^f/f^ Vasa-Cre−control females. The fathers in both cases are WT. A significant difference between Cre+ and Cre−female fertility is observed (p<0.0001, Mann-Whitney test). **d)** H&E staining of ovaries, showing normal morphology in Mettl3^f/f^ Zp3-Cre+ ovaries. **e)** Number of pups per plug produced by mating a Mettl3^f/f^ Zp3-Cre+ female, compared to a Mettl3^f/+^ Zp3-Cre+ control female. The fathers in both cases are WT. A significant difference between f/f and f/+ female fertility is observed (p<0.0001, Mann-Whitney test). **f)** Number of oocytes per mouse produced by mating Mettl3^f/f^ Zp3-Cre+ females, compared to Mettl3^f/f^ Zp3-Cre−control females. The fathers in both cases are WT. A significant difference between the number of oocytes of f/f Cre+ and f/f Cre−is observed (p<0.0002, Mann-Whitney test). **g)** Top: Experimental design - Mett3^f/f^ Zp3-Cre+ and Cre−as control underwent hormone priming, flush, fixation and staining for tubulin. Bottom: Number of oocytes observed in the different stages of meiosis. In the control, most of the oocytes were in MI stage, in KO (Cre+) all of the observed oocytes were in the GV state. **h)** Staining examples of oocytes in the different stages of meiosis as observed in KO (Cre+) and control (Cre−). **i)** Immunostaining of Ythdf1, Ythdf2 and Ythdf3 in ICR WT oocytes after hormone priming (PMS & hCG).

Next, we tested the role of Mettl3 in spermatogenesis. Mettl3^f/f^Vasa-cre+ male mice, in which the KO was activated during primordial germ cells, showed a massive reduction in the testis volume (**Figure 2a**), severe degenerative defects (**Figure S4a),** and sterility (**Figure 2b**), as was reported elsewhere (Xu et al. 2017; Lin et al. 2017). Similarly, a dissection of Mettl3^f/f^Stra8-cre+ male mice, in which the KO was activated during early-stage spermatogonia, showed a significantly reduced testis volume **(Figure 2c)**, mild degenerative changes in seminiferous tubules (**Figure S4b**), and ~75% reduction in sperm quantity observed in the cauda epididymis (**Figure S4b**). Indeed, Mettl3^f/f^Stra8-cre+ mice showed significant hypofertility compared to their counterpart control (**Figure 2d**), similar to what was previously reported (Lin et al. 2017). Interestingly, Mettl3^f/f^Prm1-cre+ male mice, in which the KO was activated in the spermatids (**Figure 2e**), showed normal fertility (**Figure 2f**) and typical seminiferous tubules morphology (**Figure S4c**).

**Figure 2.**
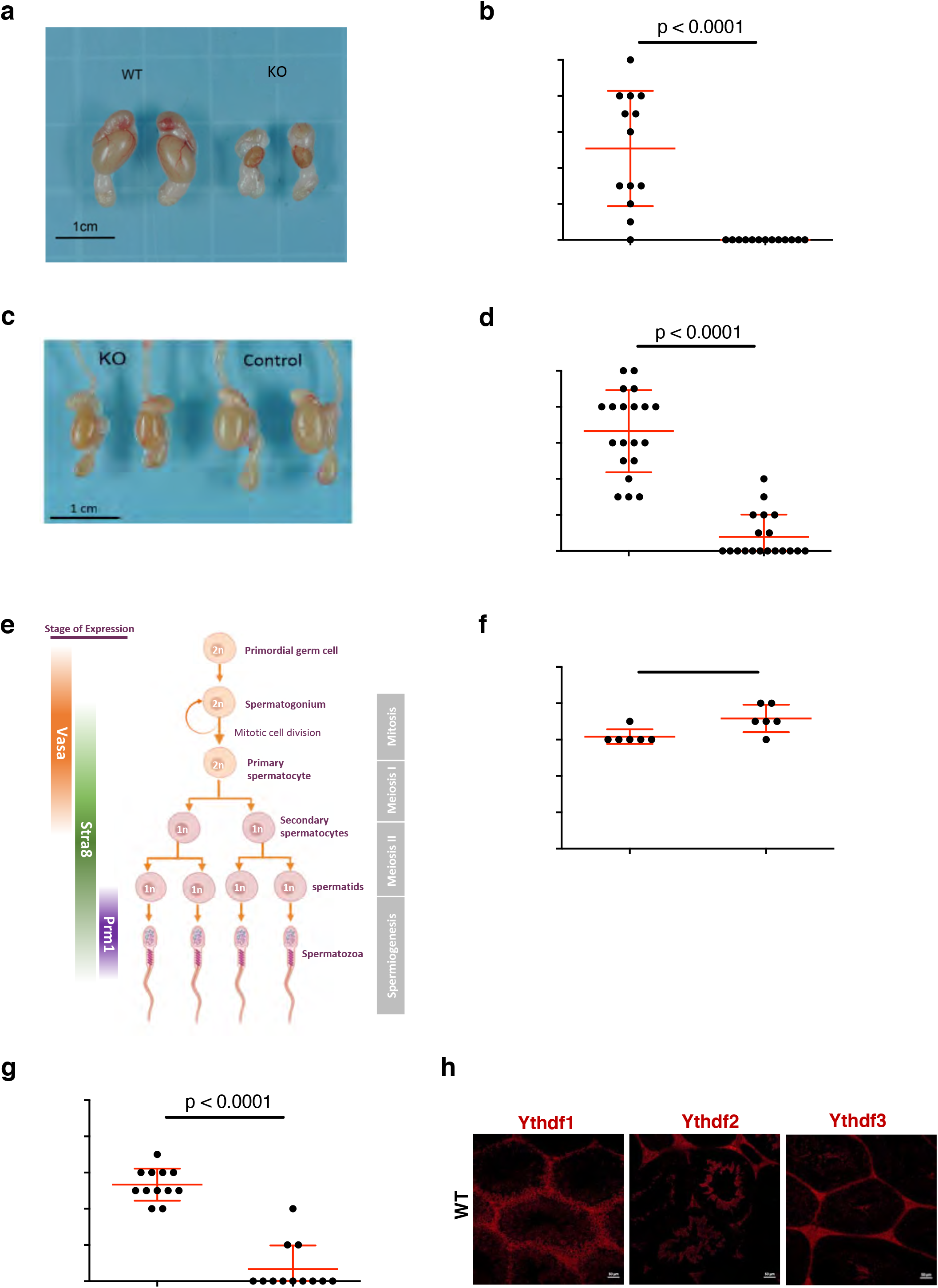
Mettl3 and Ythdf2 are essential for male mice fertility. **a)** Gross morphology of testis and epididymis of Mettl3f/f Vasa-Cre+ and Mettl3f/f Vasa-Cre−males. Cre+ males show a massive decrease in testis and epididymis size compared to Cre−control. **b)** Number of pups per plug produced by Mettl3f/f Vasa-Cre+ males, compared to Mettl3f/f Vasa-Cre−control males. The mothers in both cases were WT. In this case there is a significant hypofertility of the KO (p<0.0001, Mann-Whitney test). **c)** Gross morphology of testis and epididymis of Mettl3f/f Stra8-Cre+ and Mettl3f/f Stra8-Cre−males. Cre+ males show a reduced-size testis and epididymis compared to Cre−control. **d)** Same as in (b), for Stra8-Cre, showing a significant hypofertility of the KO (p<0.0001, Mann-Whitney test). **e)** Vasa, Stra8 and Prm1 are expressed during spermatogenesis, in different stages, as indicated. **f)** Same as in (b), for Prm1-Cre, showing no significant difference between Cre+ and Cre−male fertility. **g)** Number of pups per plug produced by Ythdf2^−/−^ males, compared to Ythdf2^+/−^ control males. The mothers in both cases were WT. A significant difference between the fertility of KO and heterozygous males is observed (p<0.0001, Mann-Whitney test). **h)** Immunostaining of Ythdf1, Ythdf2 and Ythdf3 in seminiferous tubules, showing that each of the proteins is expressed at different stages of spermatogenesis.

In summary, our genetic dissection of Mettl3’s role during gametogenesis shows the pivotal role of m^6^A modifications in both oogenesis and spermatogenesis. The severity of the phenotype was dependent on the stage in which Mettl3 was depleted. Therefore, we tested the effect of Ythdf KO, which might have a milder effect on the process.

### Ythdf2 is the only Ythdf reader which is essential for gametogenesis, and has a different expression pattern than Ythdf1 and Ythdf3

To understand the role of the Ythdf family in gametogenesis, we knocked out each of the three readers using CRISPR-Cas9 (**Figure S1c**). Heterozygous mice were further crossed to give full KO for each of the Ythdfs. The mice that were born were validated by genotyping (**Figure S1d**). Ythdf1^−/−^ and Ythdf3^−/−^ mice were born in the expected Mendelian ratio (**Figure 3a**) and showed no apparent defects. However, 80-83% of Ythdf2^−/−^ pups died shortly after birth, leading to a sub-Mendelian ratio 30 days after birth (**Figures 3a**).

**Figure 3.**
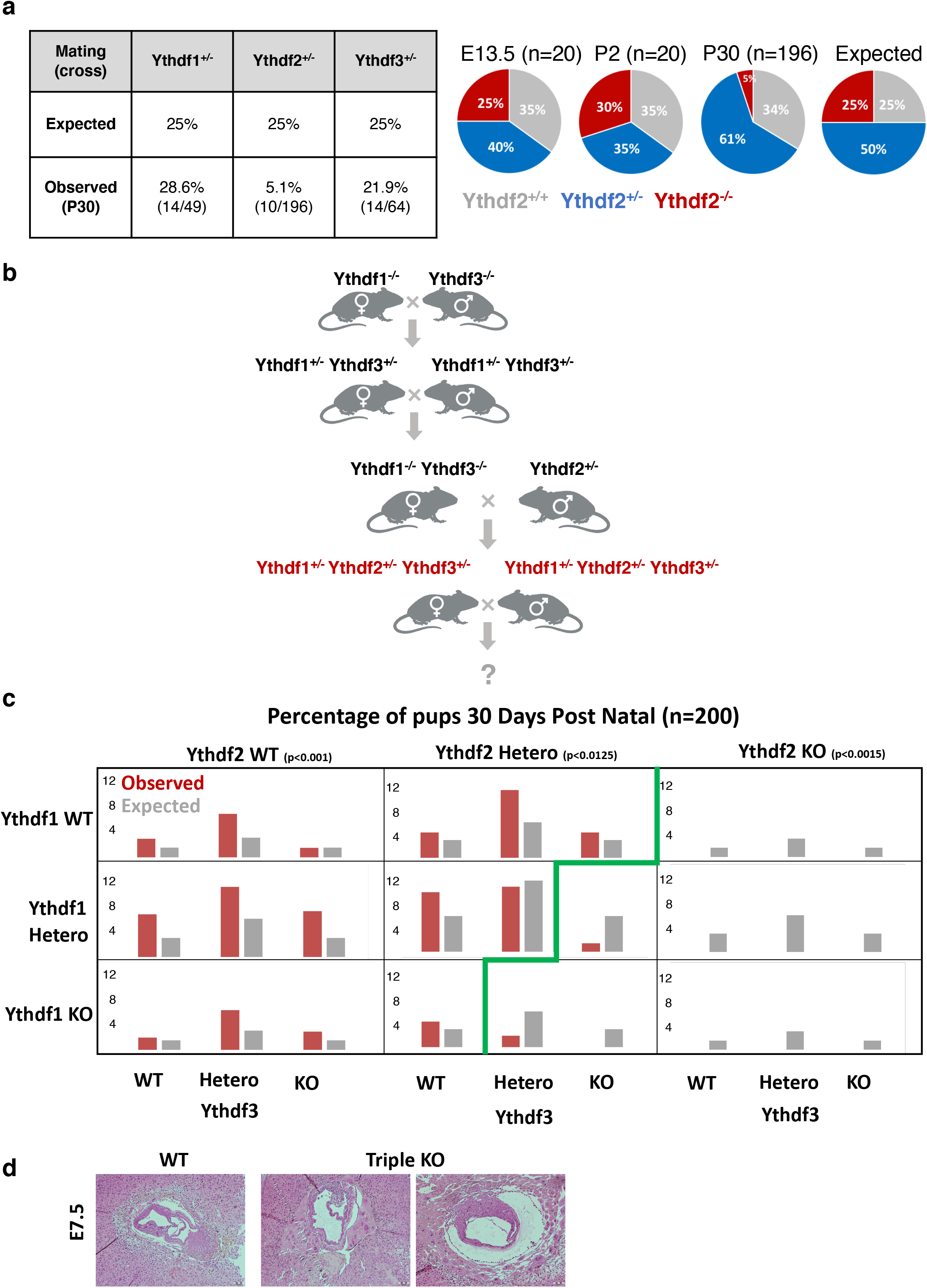
Characterization of Ythdf1-KO, Ythdf2-KO and Ythdf3-KO mice. **a)** Left: statistics of KO offspring received from crossing heterozygous mice from each of the indicated strains (Ythdf1^+/−^, Ythdf2^+/−^, Ythdf3^+/−^). Right: distribution of Ythdf2 WT, HET and KO offspring in E13.5, two days postnatal (DPN), and 30 DPN (compared to expected ratios). **b)** Crossing strategy for generating triple-heterozygous mice, which were further crossed, and their offspring, statistics are presented in panel (c). **c)** Percentage of genotypes received by crossing triple-heterozygous mice, out of 200 pups tested 30 DPN. Red - observed percentage, grey - expected under null assumption. No pups with Ythdf2-KO genotype survived 30 DPN. In the Ythdf2^+/−^ genotype, pups with KO in either Ythdf1 or Ythdf3 were born at a sub-Mendelian ratio. Chi-square test p-values are indicated. **d)** H&E staining showing the impaired morphology of triple-KO E7.5 embryos, compared to WT control.

Ythdf1-KO and Ythdf3-KO mice were fertile as their control counterparts **(Figure S5a-d)** and did not show any histological defect in their reproductive organs **(Figure S5e-f)**. As for Ythdf2, the few viable KO mice that survived were tested. While Ythdf2-KO female mice showed normal oocyte morphology **(Figure S6a)**, they were sterile **(Figure S6b)**, suggesting a later defect than what was observed in Mettl3^f/f^Zp3-cre+ oocytes. Indeed, a recent work showed that Ythdf2-KO oocytes could be fertilized but do not develop beyond the 8-cell stage, probably because of their inability to reduce maternal RNA (Ivanova et al. 2017). Next, the oocytes were flushed after hormone priming and measured for RNA levels using SMART-seq. Although the morphology was indistinguishable from WT, on the molecular level, the cells of Ythdf2-KO were already distinctly clustered compared to WT **(Figure S6c)**. Among the 311 genes that were downregulated in the KO **(Table S1)**, 72 are related to extracellular matrix (p<1.35e-06), which is crucial for oocyte fertility. Interestingly, in sperm too, m^6^A regulation of metallopeptidase is mediated by Ythdf2 (Huang et al. 2020).

Ythdf2-KO male mice showed mild degenerative changes in the seminiferous tubules (**Figure S4d**), and severe loss of sperm in the cauda epididymis (**Figure S4e**). Accordingly, these males were hypofertile (**Figure 2g**). Measuring expression in WT and KO round sperm cells, we observed changes in expression in 301 genes (**Figure S4f, Table S1**), many of them associated with cytoskeleton, microtubules and cilium functions **(Figure S4f)**, possibly explaining the impaired sperm maturation. In addition, some metallopeptidases were up-regulated in the KO (e.g. Adam4, Adamts3, Cpxm1).

Although Ythdf1 and Ythdf3 share high sequence homology with Ythdf2, they cannot compensate for its depletion during gametogenesis. One possible explanation for this observation is that the proteins are not expressed in the same spatial or temporal space during the process. To test this hypothesis, we immunostained histological sections of the testis for different reader expression. Indeed, immunostaining of seminiferous tubules showed that Ythdf1, Ythdf2 and Ythdf3 are expressed in different cells during the sperm maturation process (**Figure 2h**). Similarly, staining of GV oocytes showed a different expression pattern for Ythdf2, as it is the only Ythdf reader that is located both in the cytoplasm and nucleus **(Figures 1i, S7**). Thus, the different expression pattern of the Ythdf readers in gametogenesis may explain the lack of compensation for Ythdf2 depletion.

### Ythdf readers compensate one another in a dosage-dependent manner in early development

Next, we tried to generate Ythdf triple-KO mice for a more comprehensive understanding of the readers’ roles *in-vivo*. Ythdf1^−/−^ and Ythdf3^−/−^ mice were crossed, and the double heterozygote offspring were further crossed with Ythdf2^+/−^ mice to generate triple heterozygote mice to all three readers (Ythdf1^+/−^Ythdf2^+/−^Ythdf3^+/−^ or “triple-HET”) (**Figure 3b**). Triple-HET mice were crossed, and all their offspring (n=200) were genotyped on day 30 post-natal (**Figure 3c**). The ratio of offspring with Ythdf2-WT genotype was as or above the expected Mendelian-ratio, while no offspring with Ythdf2-KO were detected. Interestingly, even when the mice were only heterozygote for Ythdf2 (Ythdf2^+/−^), the Mendelian-ratio of offspring with another missing reader was below the expected ratio (Ythdf1^−/−^2^+/−^3^+/−^, threefold below expected, Ythdf1^^+/−^^2^+/−^3^−/−^, fourfold below expected, p<0.012). Moreover, we could not detect any offspring which were Ythdf2^+/−^ and null in the two other readers (Ythdf1^−/−^Ythdf2^+/−^Ythdf3^−/−^). These results suggest that lack of Ythdf1 or Ythdf3 can be compensated by the two other readers. However, the lack of Ythdf2 cannot be compensated by Ythdf1 or Ythdf3. In addition, the fact that partial expression of Ythdf2 in the heterozygous lineage requires the expression of at least one other reader, suggests that the function of the readers is dosage-dependent, and that a certain threshold of Ythdf readers are needed to be expressed in the cell to accomplish their function.

To better understand in which stage of development the triple-KO is defected, we analyzed triple-KO embryos on embryonic day 7.5 (E7.5). We found that already in this early stage of the post-implantation development, the triple-KO embryos were broadly deformed compared to the WT control **(Figure 3d)**.

### Triple-KO mESCs show normal self-renewal ability, but have an impaired ability to differentiate

We found that the Ythdf proteins can compensate one another *in-vivo* when expressed in the same cells, and that this compensation is dosage-dependent. Next, we wanted to better understand the molecular mechanism in which the different Ythdfs process mRNA molecules and thus affect cell viability and differentiation potential. We hypothesized that mESCs would be a good model for studying the molecular role of the readers *in-vitro*, since this is a system in which we can systematically perturb the cells and test the stem-cell activity outcomes (self-renewal and differentiation). In addition, in contrast to gametogenesis, all of the Ythdf readers are expressed in the same cells (**Figure 4a**) thus enabling us to test different compensation mechanisms. We stained for the readers in mouse ESCs, and indeed all were found to be expressed in the cytosolic compartment of the cells (**Figures 4a**). Next, we knocked out each of the Ythdfs in mouse ESCs using the CRISPR/Cas9 strategy (**Figure S8a**). In addition, we generated a triple-KO line, Ythdf1/2/3 KO using a sequential CRISPR KO. All KO cell lines were validated by immuno-staining, DNA sequencing and RNA sequencing (**Figures S8–10**). □

**Figure 4.**
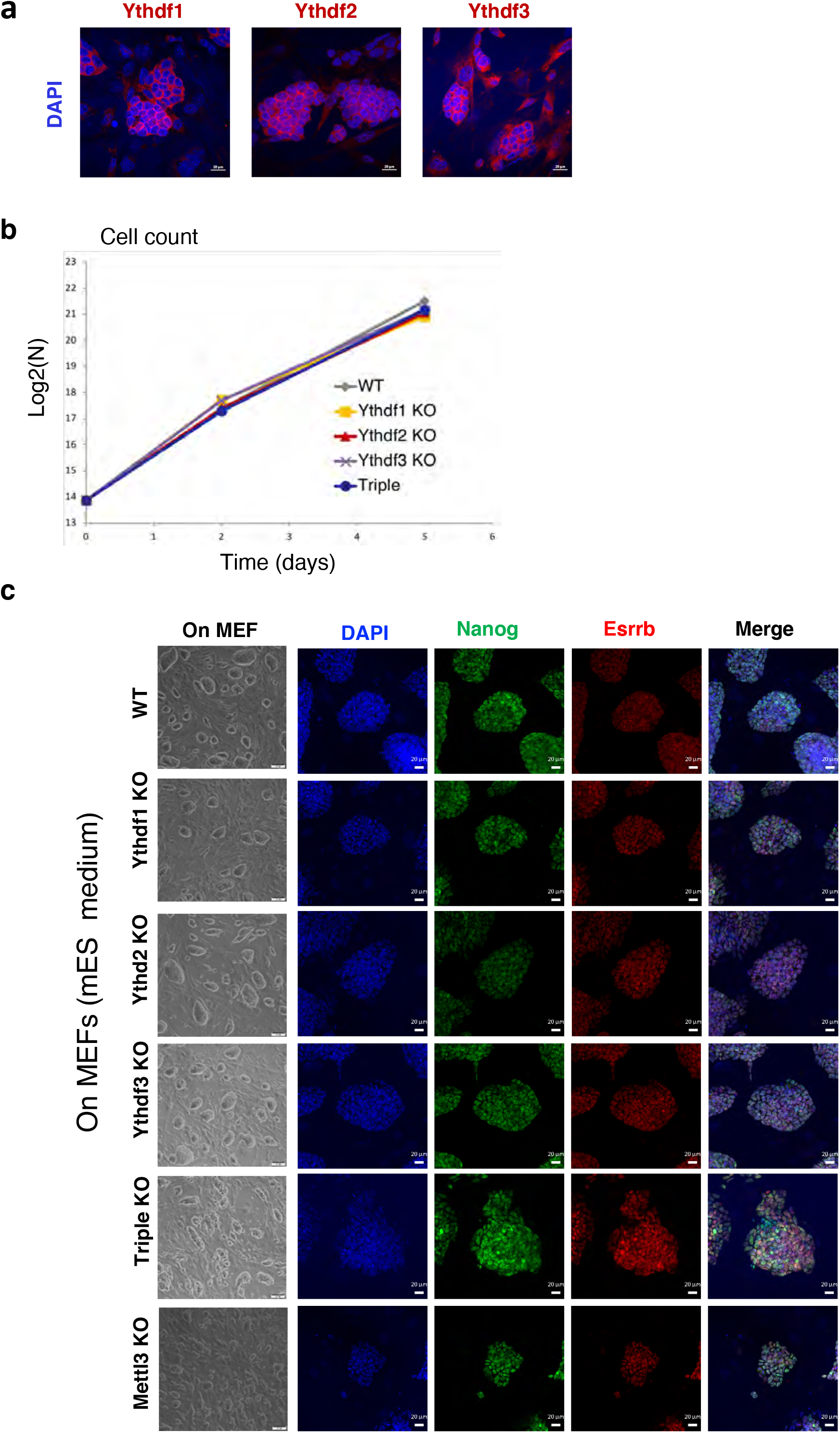
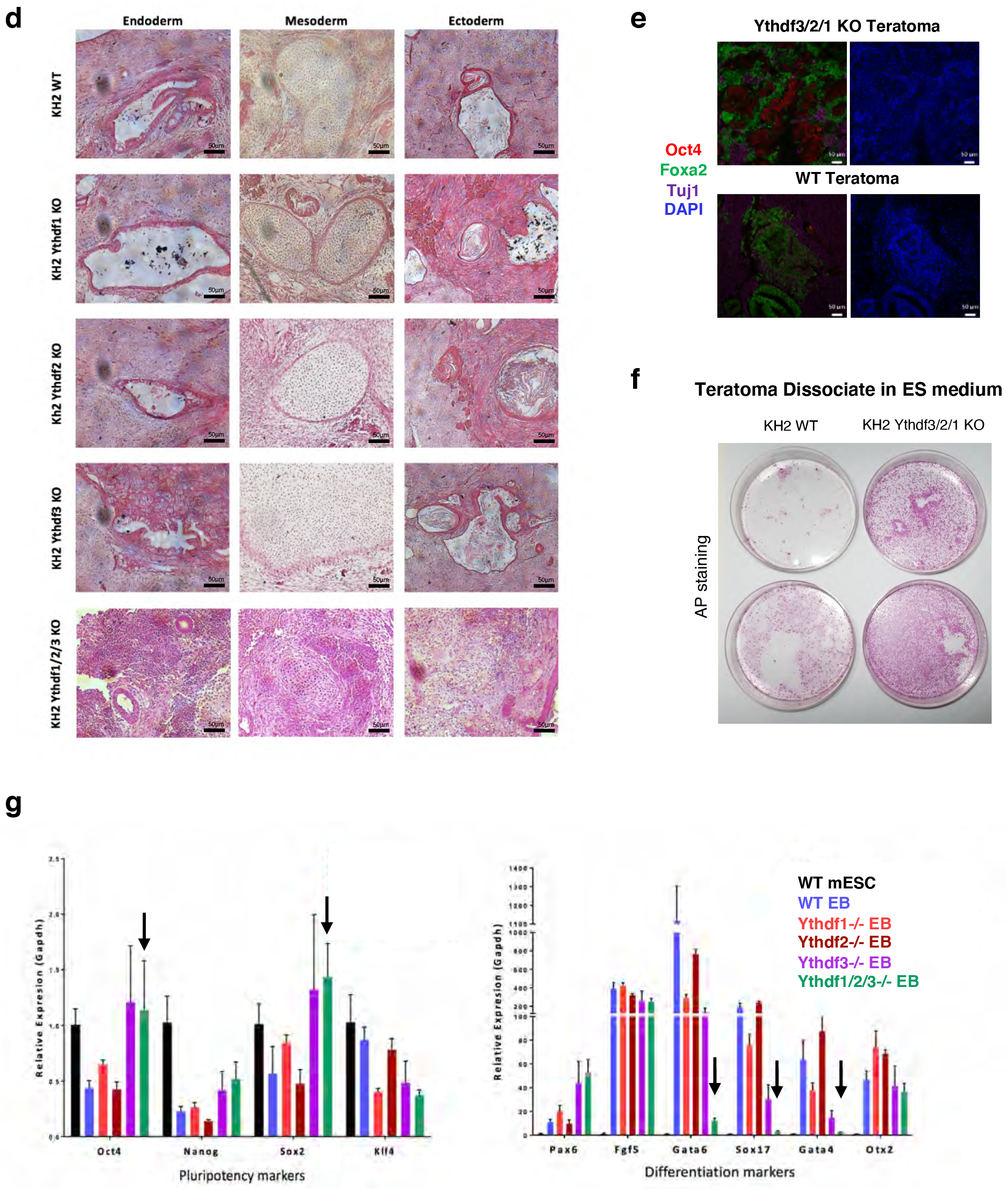
Ythdf1, Ythdf2 and Ythdf3 are redundant in ESCs differentiation. **a)** Immunostaining of Ythdf1, Ythdf2, and Ythdf3 in KH2 mESCs, showing a protein expression in the cytosolic compartment of the cell. **b)** Cell growth curve of all KO lines and WT control. Cells were grown on mouse feeders, in serum/LIF conditions. **c)** Brightfield and immunostaining of Nanog (green), Esrrb (red) and DAPI (blue), in KO cells (single, triple and Mettl3) and WT control, showing that all cell lines express Nanog and Esrrb. **d)** Teratomas generated by the KO cell line and by WT control. Single-KO cell lines show all germ layers, while triple-KO teratomas as poorly differentiated. **e)** Immunostaining of triple-KO and WT control with Oct4 (red), Foxa (green), Tuj1 (purple) and DAPI (blue). Triple-KO contains patches of Oct4 staining, unlike WT teratomas. **f)** Alkaline phosphatase (AP) staining of disassociated teratomas from Triple-KO and WT control, showing a greater AP staining in the triple-KO cells. **g)** RT-PCR of pluripotent genes (left) and differentiation genes (right), measured in WT and KO EBs, and in WT mESCs as a control. In the triple-KO EBs, pluripotent markers are higher than the control and differentiation markers are lower than the WT control.

All cell lines were viable and showed similar self-renewal ability as WT cells in mESC naïve growing conditions (**Figure 4b**). In addition all cell lines expressed normal pluripotent markers (**Figure 4c**). We next wanted to test their ability to undergo differentiation using *in-vivo* and *in-vitro assays.* First we tested their ability to generate teratomas upon injection to immune-deficient mice **(Figures 4d,e**). Interestingly, while WT and single-KO teratomas generated differentiated structures containing the three germ layers, and stained for developmental markers such as Foxa2 and Tuj1, triple-KO teratomas were poorly differentiated, and broadly stained for Oct4, a pluripotent marker (**Figures 4d,e**), indicating their poor ability to differentiate. To further examine the differentiation potential of our cells, the teratomas were disaggregated and cultured in mouse ESC medium for six days. Triple-KO cells generated significantly more pluripotent colonies, as shown by alkaline phosphatase staining (**Figure 4f**). In addition, embryoid bodies (EBs) were generated from all our cell lines, followed by RNA extraction and qPCR. Once again, differentiation markers were modestly expressed in the triple-KO EBs (**Figure 4g**), indicating their poor differentiation, compared to WT and single-KO cell lines. This phenotype is highly similar to the “hyper-pluripotency” phenotype which we observed previously in Mettl3-KO cells (Geula et al. 2015).

### The transcriptional profile of triple-KO is distinct from WT and single-KO

To dissect the molecular profile of single-KOs and the triple-KO, transcription profiles were measured using RNA-seq from each of the cell lines (Ythdf1^−/−^, Ythdf2^−/−^, Ythdf3^−/−^, Ythdf1/2/3^−/−^). In addition, we had a WT control and a positive control consisting of cells that are knocked-out to Mettl3, which lack m^6^A methylation and were previously shown to be hyper-pluripotent (Geula et al. 2015). Clustering the samples based on their transcriptional profile showed that while single-reader-KO samples cluster together with WT samples, triple-KO samples clusters more closely to Mettl3^−/−^ samples (**Figures 5a, S11a**), suggesting that single-KOs do not have a dramatic effect on the transcription profile of the cells, supporting our redundancy hypothesis.

**Figure 5.**
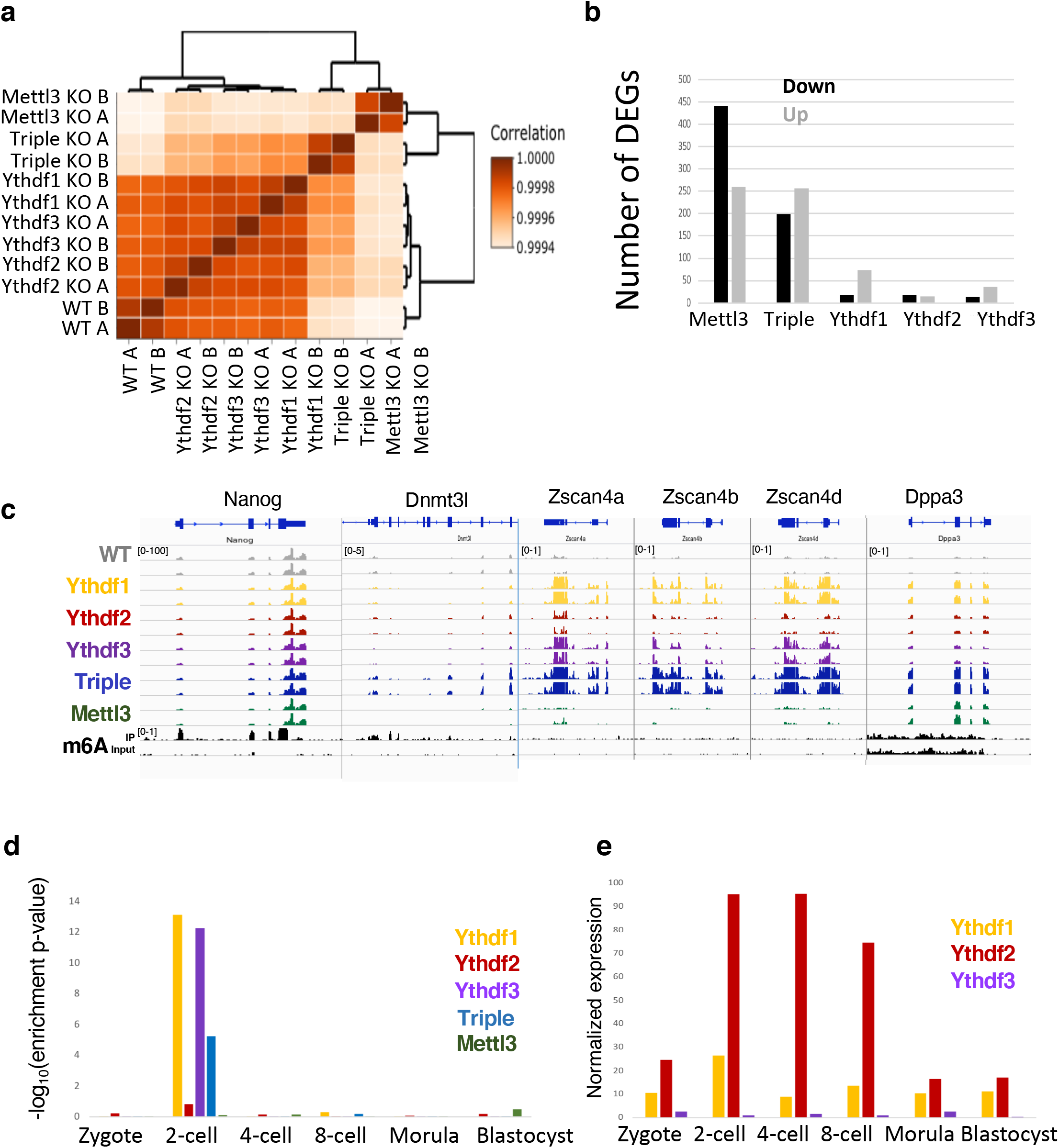
Triple-KO has a dramatic effect on gene expression. **a)** Hierarchical clustering of samples based on Pearson correlations, showing that only single-KO samples are highly similar to WT. **b)** Number of differentially expressed genes in each of the KO cell line, compared to WT. Black: downregulated genes. Grey: upregulated genes **c)** RNA-seq and m^6^A methylation landscape of selected genes. Normalized coverage is presented. Only Nanog and Dnmt3l are m^6^A-methylated. Dnmt3l, Zscan4a b d & Dppa3 are over-expressed in triple-KO. **d)** Enrichment of upregulated genes in each category, to early embryo genes (Gao et al. 2017b). Genes that are upregulated in KO of Ythdf1 & 3 are specifically enriched for two-cell genes. **e)** Normalized expression of Ythdf1,2 & 3, as measured in early mouse embryo (Gao et al. 2017b).

When we analyzed the number of differentially expressed genes in each of the cell lines (**Figure 5b, Table S2**), we could see that the few genes that were upregulated in the single-KOs (77 In Ythdf1, 16 in Ythdf2, 37 in Ythdf3), greatly overlapped with the genes that were up-regulated in triple-KO and to a lesser extent in Mettl3-KO (**Figure S11b**). Interestingly, several of the genes that were upregulated in Ythdf1-KO,Ythdf3-KO and triple-KO, but not in Ythdf2-KO, were significantly enriched for two-cell stage embryo genes (genes that are expressed after the first division of the zygote), such as Zscan4a,c,d,f, Usp17a,b,c,e, Zfp352, Gm20767 and Tcstv1 (**Figure 5c, 5d and S11b**) (Storm et al. 2009). This suggests the even though most of Ythdfs’ effects on transcription are redundant, some Ythdf readers cannot always compensate for the absence of the others.

To further investigate the role of the Ythdfs, their RNA binding target profile was measured in three different single-Ythdf flag-tagged mESC lines using the eCLIP method (Van Nostrand et al. 2016). We found 201, 1995 and 146 targets that were bound by Ythdf1, Ythdf2 and Ythdf3 respectively. All the found targets significantly overlapped with previously published data measured in human (Patil et al. 2016; Wang et al. 2015; Li et al. 2017; Niu et al. 2013; Shi et al. 2017) (**Figure S12a-c)**. In addition, a significant part of the genes bound by the readers, also carry m^6^A methylation, 75%, 70% and 63% for Ythdf1, Ythdf2 and Ythdf3 respectively **(Figure S12d)**. When we analyzed the common peaks between the readers, we found that the peaks bound by Ythdf1 and Ythdf3 highly overlap the peaks bound by Ythdf2 (72% and 49%, respectively, **Figure S12e, Table S3**), indicating again a possible redundancy between the readers’ binding sites. However, targets of Ythdf1 and Ythdf3 were not enriched for two-cell genes, which are typically not expressed in mESCs, but rather to blastocyte genes that are expressed in mESCs **(Figure 12f)**. To investigate the roles of Ythdf1&3 in the context of two-cell genes, binding profiling needs to be done in two-cell stage embryos, which is not feasible with the current technology (Van Nostrand et al. 2016).

### Significant increases in m^6^A methylated mRNA half-life seen in only the triple-KO

Previous studies proposed that m^6^A has a role in mRNA degradation (Du et al. 2016; Geula et al. 2015), specifically, Wang et al. (2014) suggested that Ythdf2 binds methylated transcripts and directs them to mRNA decay sites. We therefore examined the decay rate in mouse ESCs, in each of the single-KO and triple-KO cells. We treated the cells with actinomycin-D, and harvested RNA at three time points (t=0, 4h, 8h, with duplicates of 0 and 8). We estimated transcription levels using 3’ poly-A RNA-seq (Geula et al. 2015), and calculated the mRNA half-life **(Methods, Table S4)**. In single-KO cells, including Ythdf2-KO, m^6^A-methylated mRNA was degraded at a similar rate to non-methylated mRNA **(Figure 6a)**. Only in the triple-KO cells we observed a significant increase in the half-life of m^6^A methylated mRNA, compared to non-methylated mRNA, similar to what was observed in Mettl3-KO **(Figure 6a)**. The fact that in single-KOs there was no significant effect on degradation, suggests that all of the readers have similar roles in mRNA degradation and can compensate one another.

**Figure 6.**
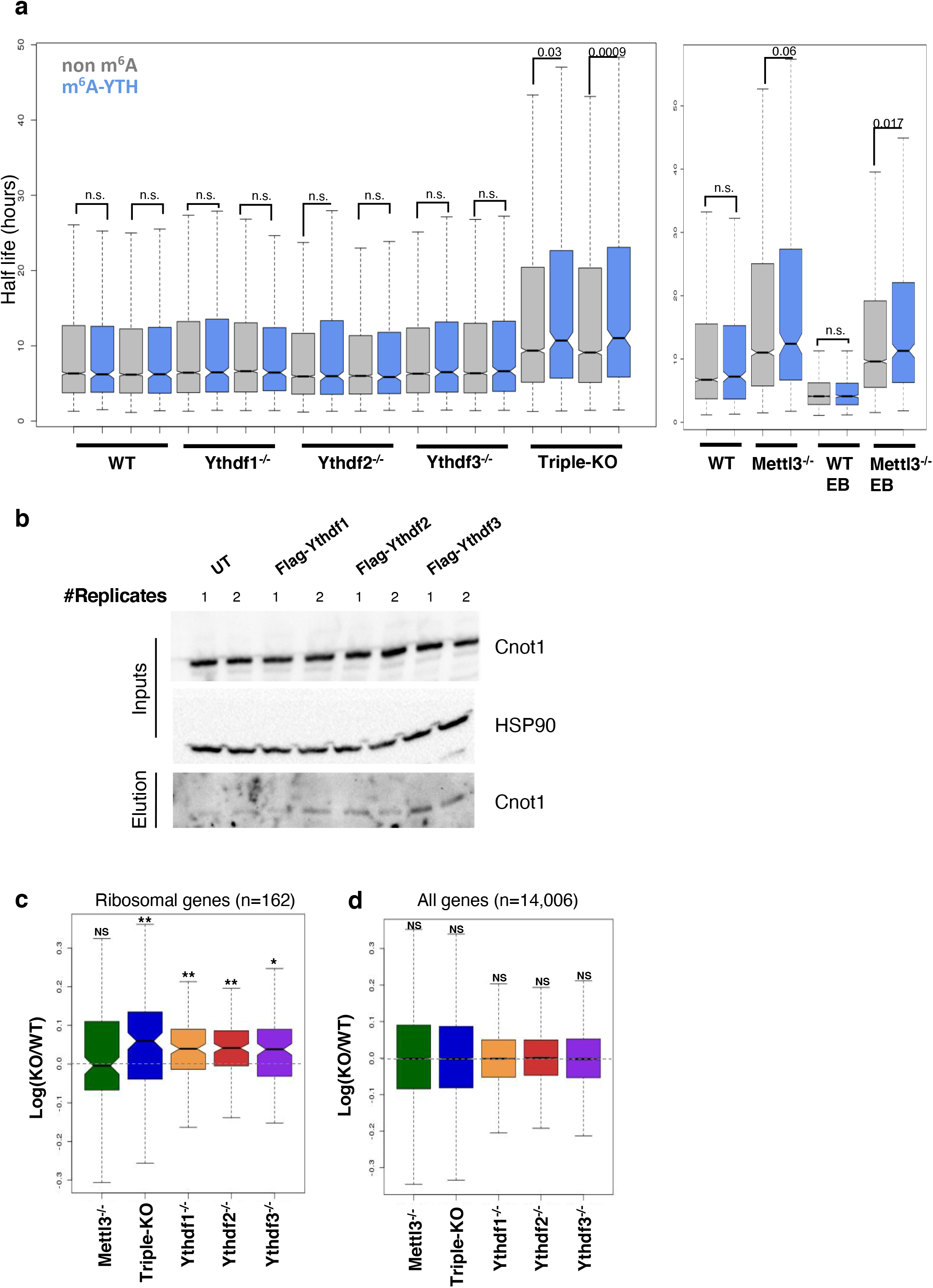
All Ythdf readers interact with Cnot1 and promote mRNA degradation. **a)** Half-life calculated in non-m^6^A genes (grey) and m^6^A-genes (blue), in each of the KO cell lines and WT control. Only in the Triple-KO and Mettl3-KO are there a significant difference between the half-life of m^6^A and non-m^6^A genes. **b)** Flag Tag coIP of CNOT1 and HSP90 as a control, showing interactions between Ythdf1 2&3 to CNOT1. c) LogRatio(KO/WT) distribution of ribosomal genes (n=162), showing a significant increase in their expression in the single and triple KO. * p-value < 10^−6^ ** p-value < 10^−10^ (paired Wilcoxon test). d) LogRatio(KO/WT) distribution of all genes (n=14,006), showing a non-significant difference in their expression (p>=0.01, paired Wilcoxon test).

A previous study (Du et al. 2016) showed that Ythdf2 recruits the CCR4-NOT complex to mediate accelerated deadenylation and decay. Interestingly, we observed that also Ythdf1 and Ythdf3 interact with CNOT1, a subunit of the CCR4-NOT complex **(Figure 6b)**. Indeed, Ythdf1 and Ythdf3 were also shown to promote deadenylation (Du et al. 2016), further supporting the hypothesis that the three readers contribute to mRNA decay, and may compensate in case of a partial loss.

Translation was also reported as a possible biological process that is affected by m^6^A methylation (Shi et al. 2017). We therefore set to measure translation in our cell lines, using a ribosomal footprint, which measures fragments of mRNA that are bound to a ribosome (Stern-Ginossar et al. 2012). To compare translation, the ribosomal footprint was normalized by mRNA levels, giving a translation efficiency level for each gene in each cell line. We observed higher translation efficiency of m^6^a-methylated genes, and of Ythdf targets, compared to non-methylated genes **(Figure S13b)**. However, this difference in translation efficiency was not affected by any of the knockouts, suggesting that translation efficiency is not mediated directly by m^6^A methylation or the Ythdfs proteins. Interestingly, a mild but significant increase in the expression of ribosomal genes **(Figure 6c)** was observed in single-KO and triple-KO cell lines, an increase which is less apparent in the global gene population **(Figure 6d)**.

## Discussion

Previous papers have suggested that each of the Ythdf readers has unique functions (Shi et al. 2017; Wang et al. 2015; Li et al. 2017; Du et al. 2016; Wang et al. 2014). We propose a different model, according to which, Ythdf readers have redundant functions to some extent, and show multiple lines of evidence supporting this model.

Using a viability assay, we observed that there is compensation between the readers, and this compensation is dosage dependent: Ythdf2 full KO or Ythdf2-heterozygotes that are also null in the two other readers, are not viable. Ythdf2 heterozygote mice need at least one functional copy of another Ythdf to escape total mortality **(Figure 3c)**. The fact that Ythdf2-KO has the most severe lethality phenotype may be due to differences in expression patterns, as seen in oogenesis and spermatogenesis **(Figures 1i, 2h)**.

The strongest evidence for Ythdf redundancy was observed in mESCs, a system in which all Ythdf readers are expressed in the cytoplasmatic compartment, thus allowing examination of our hypothesis. Indeed, in mESCs, we observed a redundancy in the function of Ythdf readers. Single-KOs were viable, had normal self-renewal ability, expressed pluripotent markers and differentiated normally upon signaling. Only in the Ythdf triple-KO did we observe an impaired differentiation ability *in-vivo* and *in-vitro*, similar to the Mettl3-KO phenotype (Geula et al. 2015)). In addition, only in the triple-KO did we observe a significant decrease in the degradation rate of m^6^A methylated transcripts, while no change was observed in the single-KOs. Redundancy is also supported by the observation that all Ythdf readers were found to bind Cnot1, part of the CCR4-NOT deadenylation complex which is a suggested degradation mechanism (Du et al. 2016).

The difference in expression patterns *in-vivo* hints to us that Ythdf readers are differentially regulated. Further experiments that induce the expression of Ythdf1/3 under the promoter of Ythdf2 can strongly support this hypothesis. Such a system can be examined in additional developmental processes such as neurogenesis or hematopoiesis. Lastly, the mechanisms that regulate Ythdf expression, such as transcription factors that bind the readers, and the reader’s response to external signals, await further investigation.

## Abbreviation used

KO: knockout
WT: wild type
mESCs: mouse embryonic stem cells
EBs: embryoid bodies
GV: germinal vesicle
TE: translation efficiency

## Acknowledgments

JHH and NN are funded by Nella and Leon Benoziyo Center for Neurological Diseases; David and Fela Shapell Family Center for Genetic Disorders Research; Kekst Family Institute for Medical Genetics; Helen and Martin Kimmel Institute for Stem Cell Research; Flight Attendant Medical Research Council (FAMRI); Dr. Barry Sherman Center for Medicinal Chemistry; Pascal and Ilana Mantoux; Dr. Beth Rom-Rymer Stem Cell Research Fund; Edmond de Rothschild Foundations; Zantker Charitable Foundation; Estate of Zvia Zeroni; European Research Council (ERC-CoG); Israel Science Foundation (ISF); Minerva; Israel Cancer Research Fund (ICRF) and BSF. **We thank Schraga Schwartz, Igor Ulitsky, Tsviya Olender, Eli Arama and Aharon Nachshon for insightful discussions and support.** We thank the Weizmann Institute management and board for providing critical financial and infrastructural support.

## Author Contributions

L.L. and J.H.H. conceived the idea for this project, designed and conducted the experiments. L.L. and N.N. wrote the manuscript with J.H.H. N.N. supervised all bioinformatics analysis and analyzed the data. V.K. assisted in libraries preparation, immunostaining and tissue culture. L.L., S.G. and V.K. engineered cell lines and mice strains under S.V.’s supervision. S.G. assisted in teratoma formation, immunostaining and Western Blots. M.Z. assisted in mouse dissection and oocyte flushing. N.M. assisted in tissue culture and Western Blots. A.A.C. assisted in oocyte staining. O.M. assisted in Ribo-seq library preparation, supervised by N.S.G; J.S., A.S. and S.A. conducted the eCLIP experiments. A.N. assisted in Ribo-seq analysis. S.S. analyzed the eCLIP data. G.W.Y. supervised the execution of the eCLIP experiments and analyses. J.H.H. and N.N. supervised executions of experiments, adequate analysis of data, and presentation of conclusions made in this paper.

## Supplementary Figure Legends

**Figure S1.**
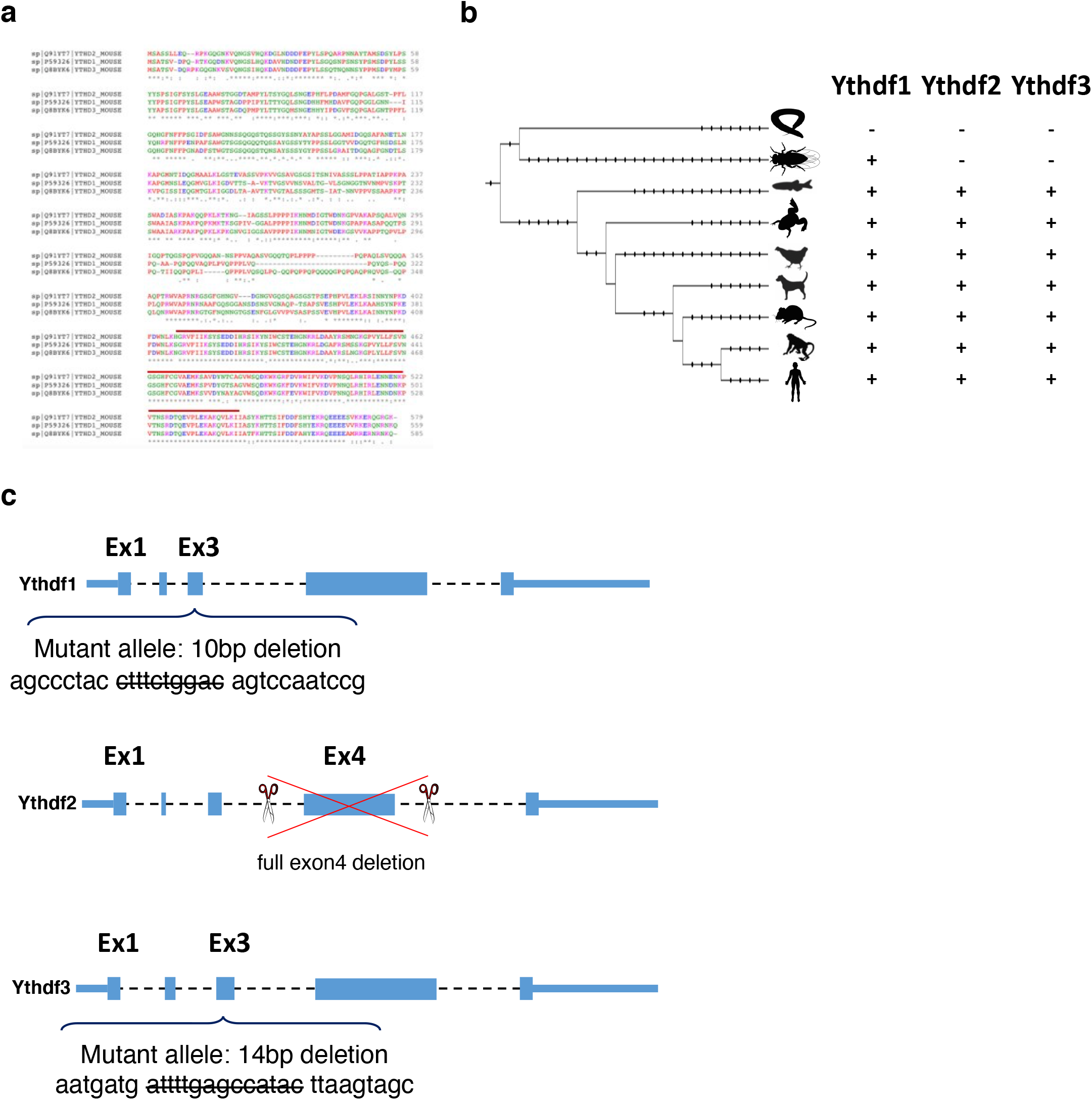
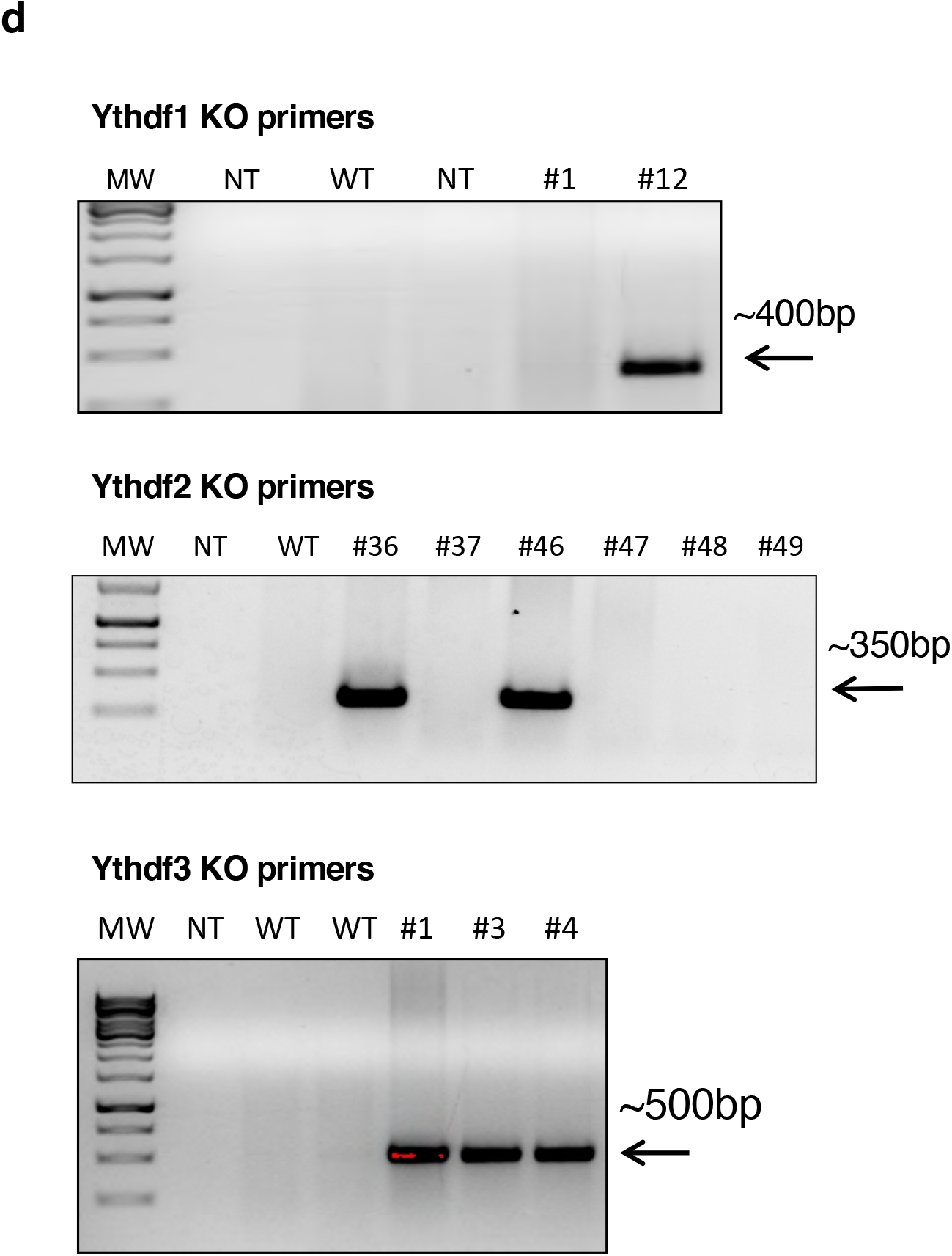
Generating Ythdf1-KO, Ythdf2-KO and Ythdf3-KO in vivo and validation. a)□ Multiple alignments of Ythdf1, Ythdf2 & Ythdf3 proteins, calculated using the Clustal Omega tool. The area of YTH-domain is highlighted in red. Ythdf1-Ythdf3 protein sequence similarity is 70.11%, Ythdf1-Ythdf2 is 67.15%, and Ythdf2-Ythdf3 is 67.78%. b)□ Phylogenetic tree of the protein sequences of Ythdf1, Ythdf2 and Ythdf3, based on the UCSC database. The three readers appear together in vertebrates, possibly due to whole genome duplication. c)□ CRISPR-Cas9 targeting strategy for knocking-out Ythdf readers in vivo. d)□ KO validation using PCR, showing successful primer integration in clones #12 (Ythdf1); #36, #46 (Ythdf2); #1, #3 & #4 (Ythdf3).

**Figure S2.**
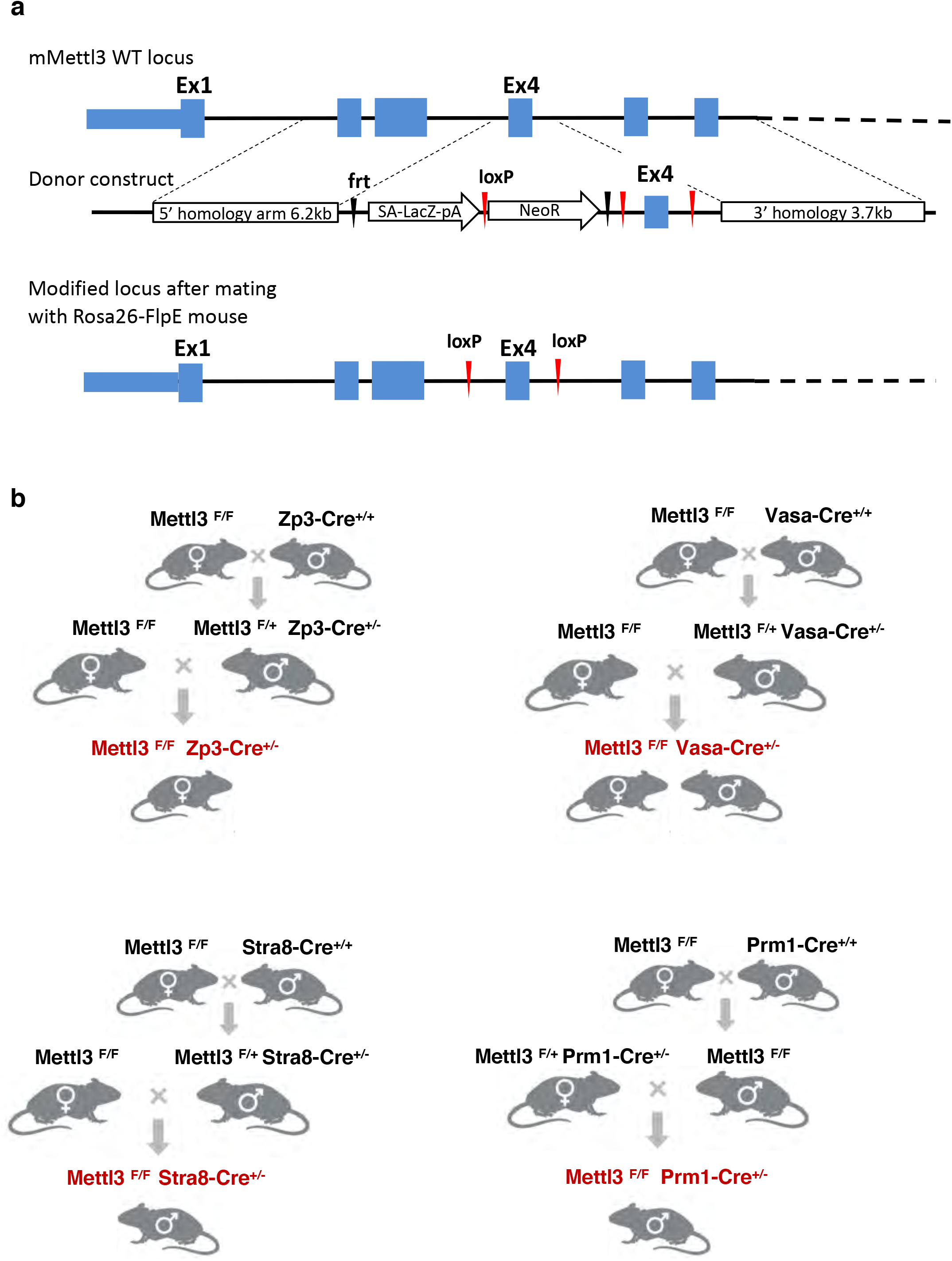
Generating conditional knockout mice models. a)□ Targeting strategy for generating Mettl3^f/f^ mice. b)□ Crossing strategy for generating different Mettl3^f/f^ Cre+ mice.

**Figure S3.**
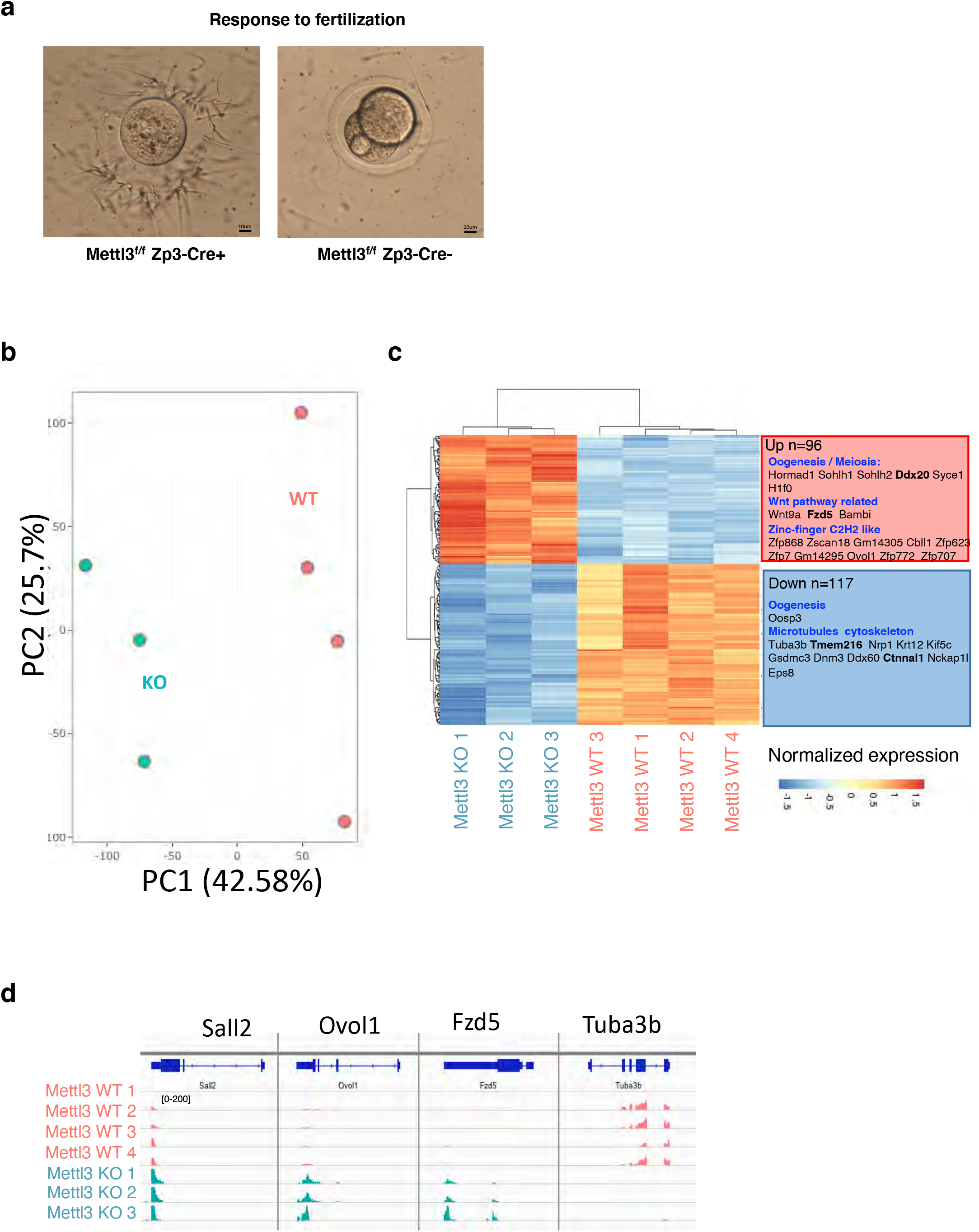
Mettl3 is essential for female mice fertility. a)□ *In vitro* fertilization of Mettl3^f/f^ Zp3-Cre−control oocytes with WT sperm, leads to creation of two-cell stage embryos, while the Mettl3^f/f^ Zp3-Cre+ oocytes fail to do so. b)□ PCA of transcriptional profile of Mettl3^f/f^ Zp3-Cre− and Mettl3^f/f^ Zp3-Cre+ oocytes, showing a distinct expression pattern. c)□ Differentially expressed genes between Mettl3^f/f^ Zp3-Cre− and Mettl3^f/f^ Zp3-Cre+ oocytes, along with selected enriched categories. m^6^A-methylated genes appear in bold. Ninety-six genes are upregulated in the KO, and 117 are downregulated in the KO. d)□ RNA-seq landscape of selected differential genes, generated with an IGV browser. Normalized coverage is presented.

**Figure S4.**
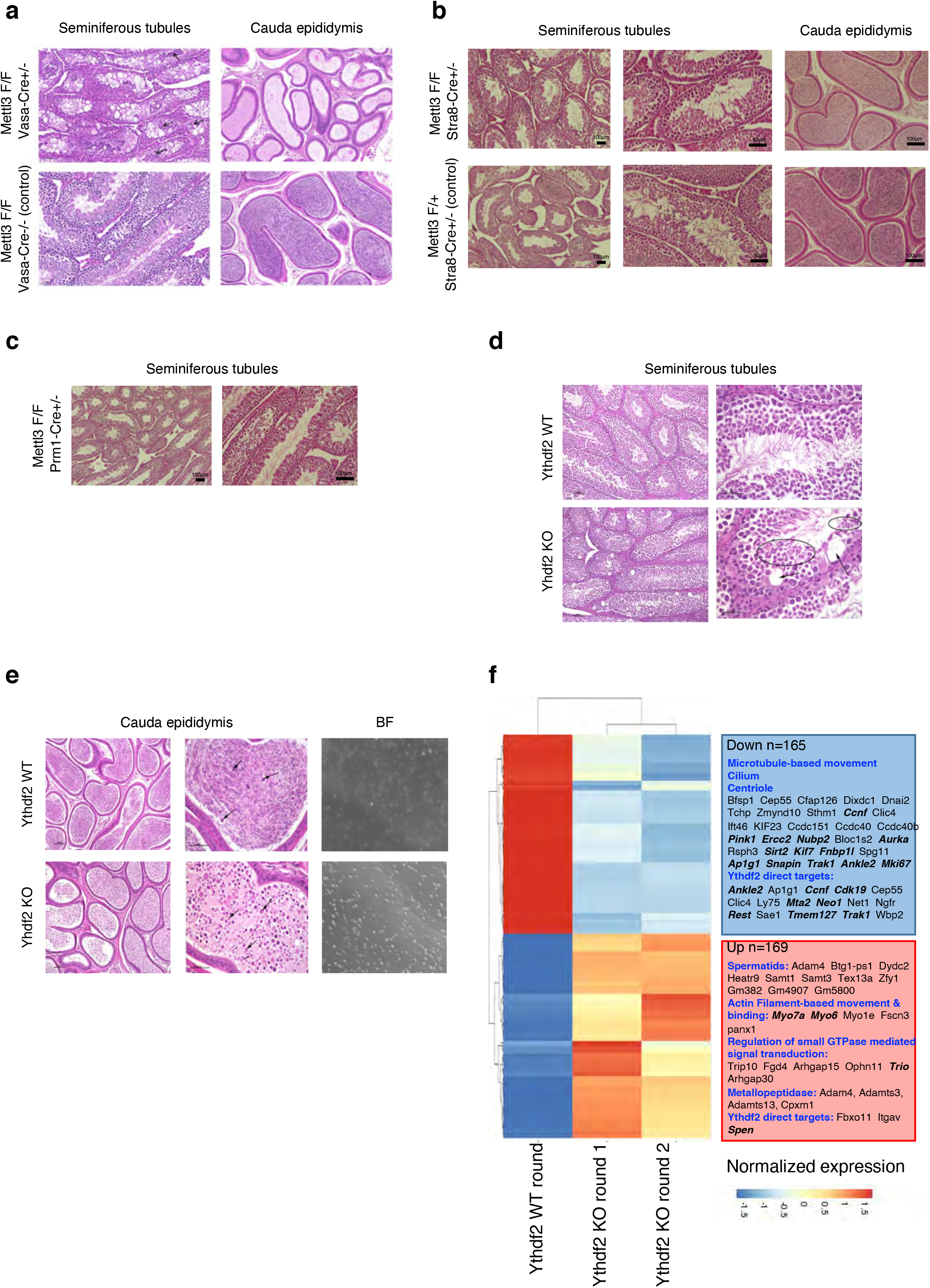
Mettl3 and Ythdf2 are essential for male mice fertility. a)□ H&E staining showing severe degenerative changes in the seminiferous tubules of Mettl3^f/f^Vasa-Cre+ and lack of sperm in the cauda epididymis. b)□ H&E staining showing mild degenerative changes in the seminiferous tubules of Mettl3^f/f^Stra8-Cre+ and ~75% reduction in sperm quantity in the cauda epididymis, compared to Mettl3^f/+^Stra8-Cre+ sibling control. c)□ H&E staining of seminiferous tubules showing a normal morphology in Mettl3^f/f^Prm1-Cre+ males. d)□ H&E staining showing mild degenerative changes in the seminiferous tubules in Ythdf2-KO males, compared to WT control. e)□ Left: H&E staining in the cauda epididymis, showing severe loss of sperm in Ythdf2-KO compared to control. Right: Brightfield of sperm extracted from the cauda epididymis of KO and control, showing a severe reduction in normal sperm quantity in the KO sample, compared to control. f)□ Transcriptional profile of genes that are differentially expressed between Ythdf2-KO and WT round spermatids, along with selected enriched categories. m^6^A-methylated genes appear in bold; 145 downregulated in KO, and 156 upregulated in KO.

**Figure S5.**
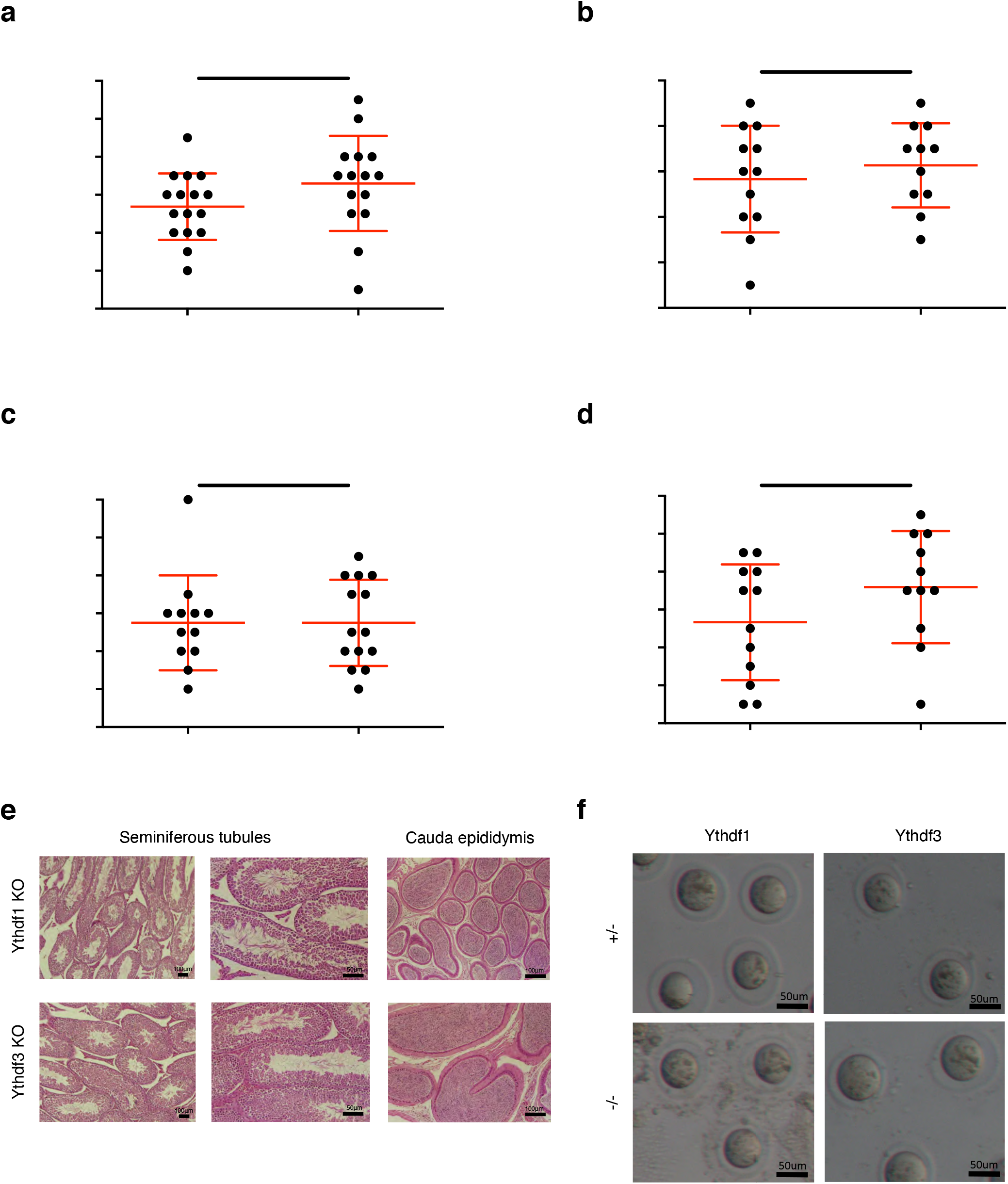
Ythdf1 knockout and Ythdf3 knockout mice are fertile. a)□ Number of pups per plug produced by mating Ythdf1-KO males, compared to Ythdf1-HET males. The mothers in both cases are WT. Here there is no significant difference between KO and HET male fertility (Mann-Whitney test). b)□ Number of pups per plug produced by mating Ythdf1-KO females, compared to Ythdf1-HET females. The fathers in both cases are WT. Here there is no significant difference between KO and HET female fertility (Mann-Whitney test). c)□ Number of pups per plug produced by mating Ythdf3-KO males, compared to Ythdf3-HET males. The mothers in both cases are WT. Here there is no significant difference between KO and HET male fertility (Mann-Whitney test). d)□ Number of pups per plug produced by mating Ythdf3-KO females, compared to Ythdf3-HET females. The fathers in both cases are WT. Here there is no significant difference between KO and HET female fertility (Mann-Whitney test). e)□ H&E staining of seminiferous tubules showing a normal morphology in Ythdf1-KO and Ythdf3-KO males. f)□ The morphology of Ythdf1-KO and Ythdf3-KO flushed oocytes appears to be normal, similar to the Ythdf1-heterozygous (HET) and Ythdf3-HET flushed oocytes.

**Figure S6.**
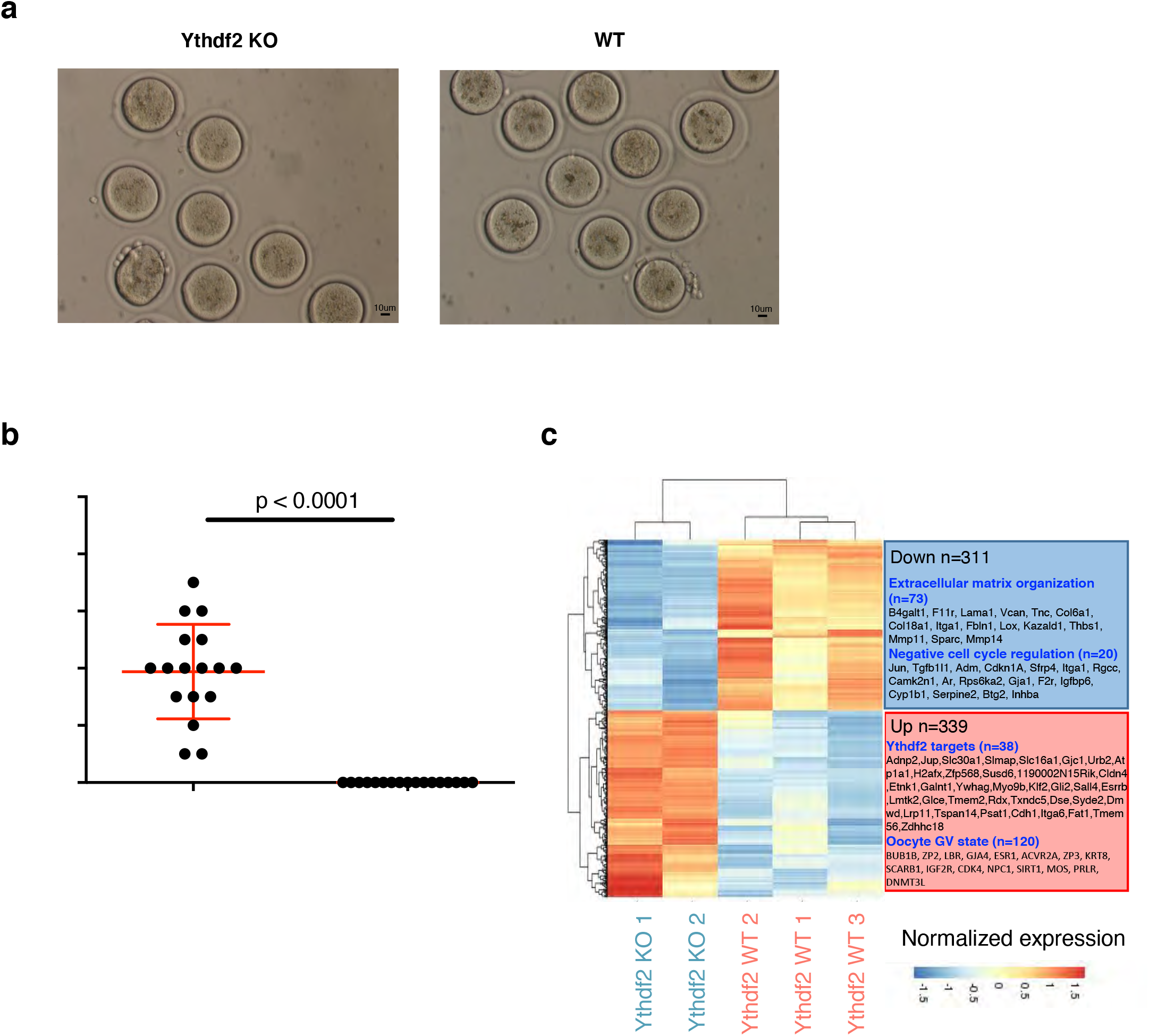
Ythdf2 is essential for female mice fertility. a)□ The morphology of Ythdf2-KO flushed oocytes appears to be normal, similar to the WT flushed oocytes. b)□ Number of pups per plug produced by mating Ythdf2^−/−^ females, compared to Ythdf2^+/−^ control females. The fathers in both cases are WT. A significant difference between the fertility of KO and heterozygous females is observed (p<0.0001, Mann-Whitney test). c)□ Transcriptional profile of genes that are differentially expressed between Ythdf2-KO and WT oocytes, along with selected enriched categories; 311 downregulated in KO, and 339 upregulated in KO.

**Figure S7.**
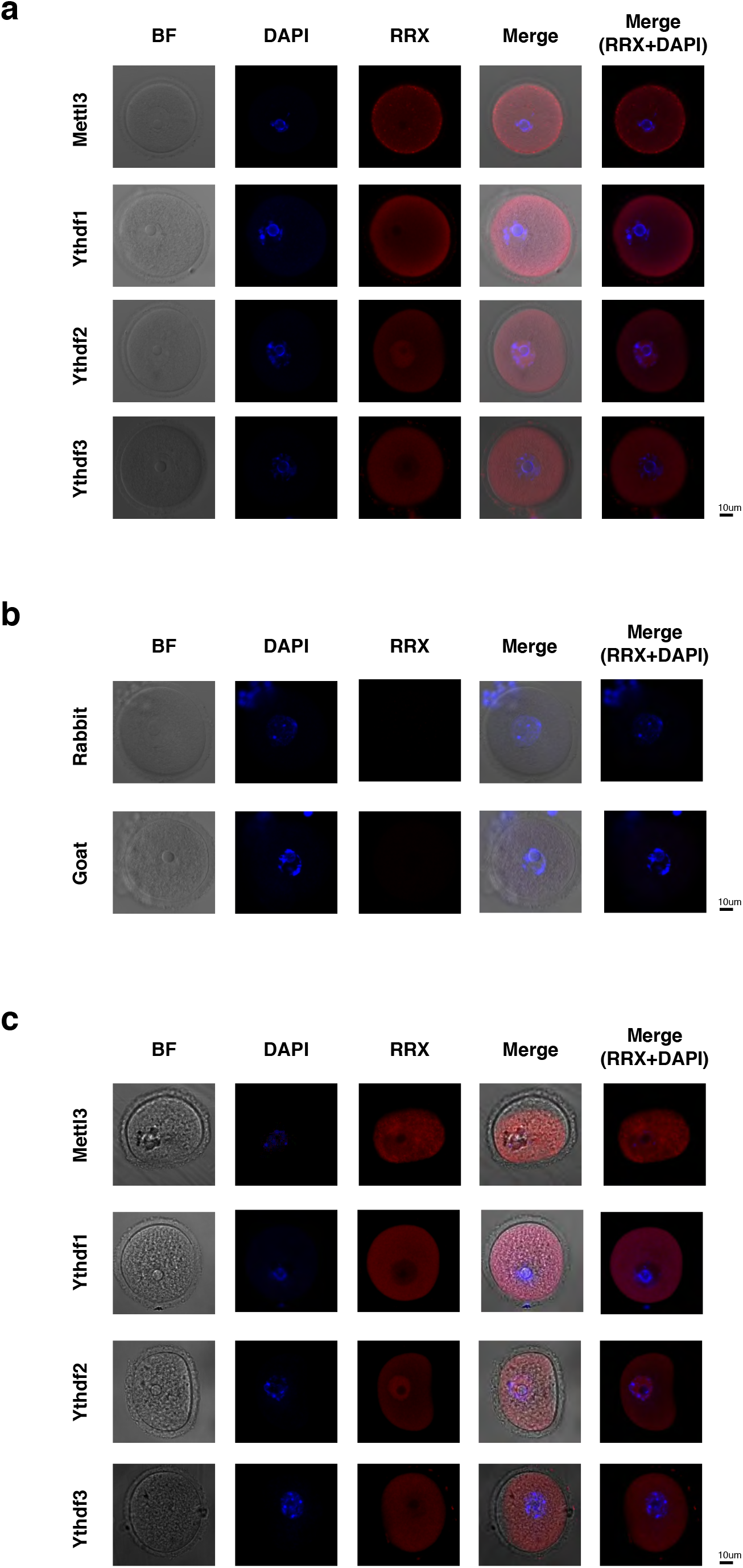
Oocytes staining for Ythdf1, Ythdf2 and Ythdf3. a)□ Immunostaining of Ythdf1, Ythdf2 and Ythdf3 in ICR WT oocytes after hormone priming (PMS & hCG). b)□ Immunostaining of Ythdf1, Ythdf2 and Ythdf3 in ICR WT oocytes after PMS & HCG - negative control (NC), without primary antibody. c)□ Immunostaining of Ythdf1, Ythdf2 and Ythdf3 in ICR WT oocytes without hormone priming.

**Figure S8.**
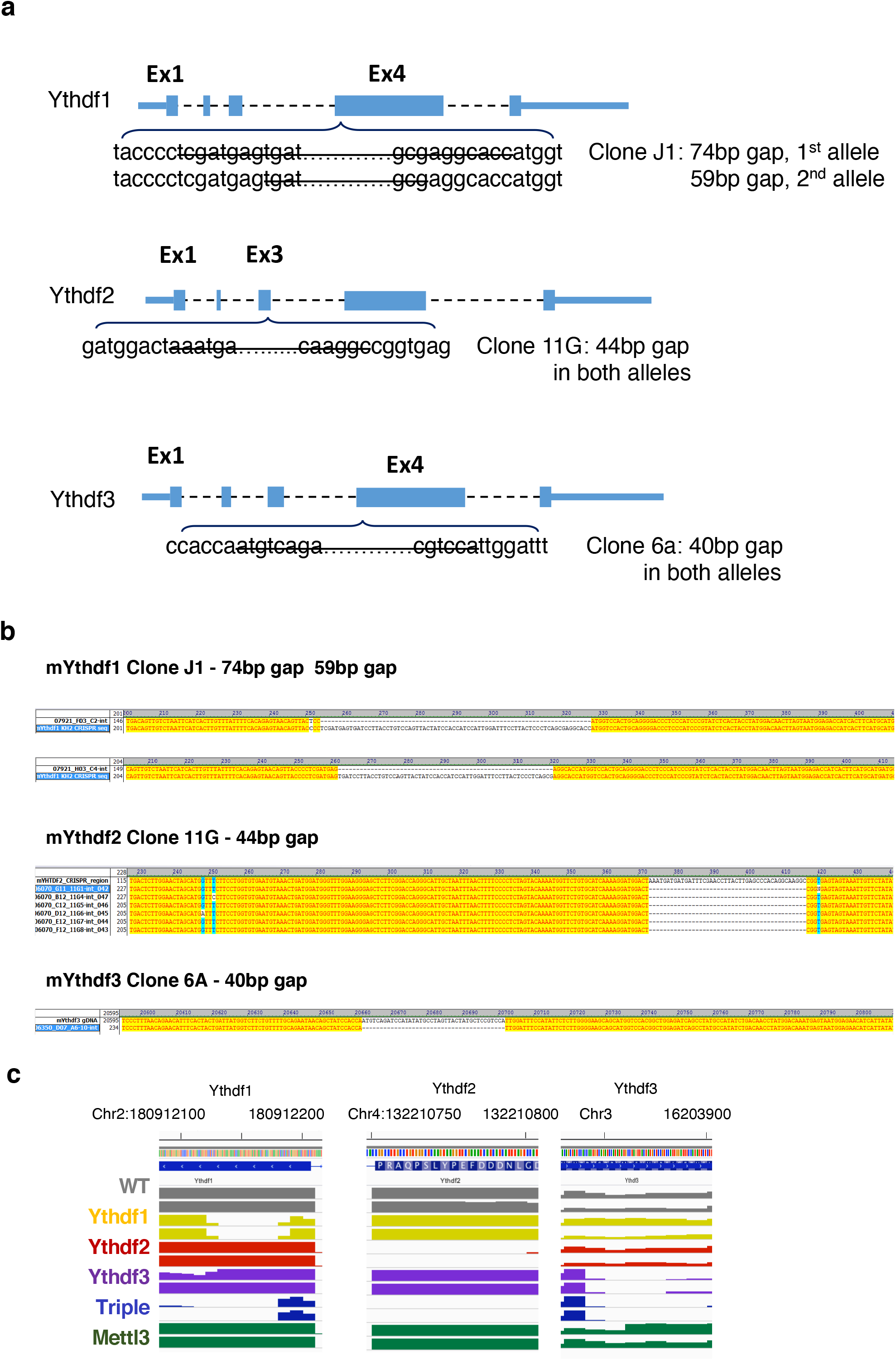
Generation and validation of knockout mESC lines. a)□ CRISPR-Cas9 targeting strategy for knocking out Ythdf readers in mESC cell lines. b)□ Sequencing validation of the single-KO lines. c)□ IGV browser view showing the missing fragments in the KO of Ythdf1, Ythdf2 & Ythdf3.

**Figure S9.**
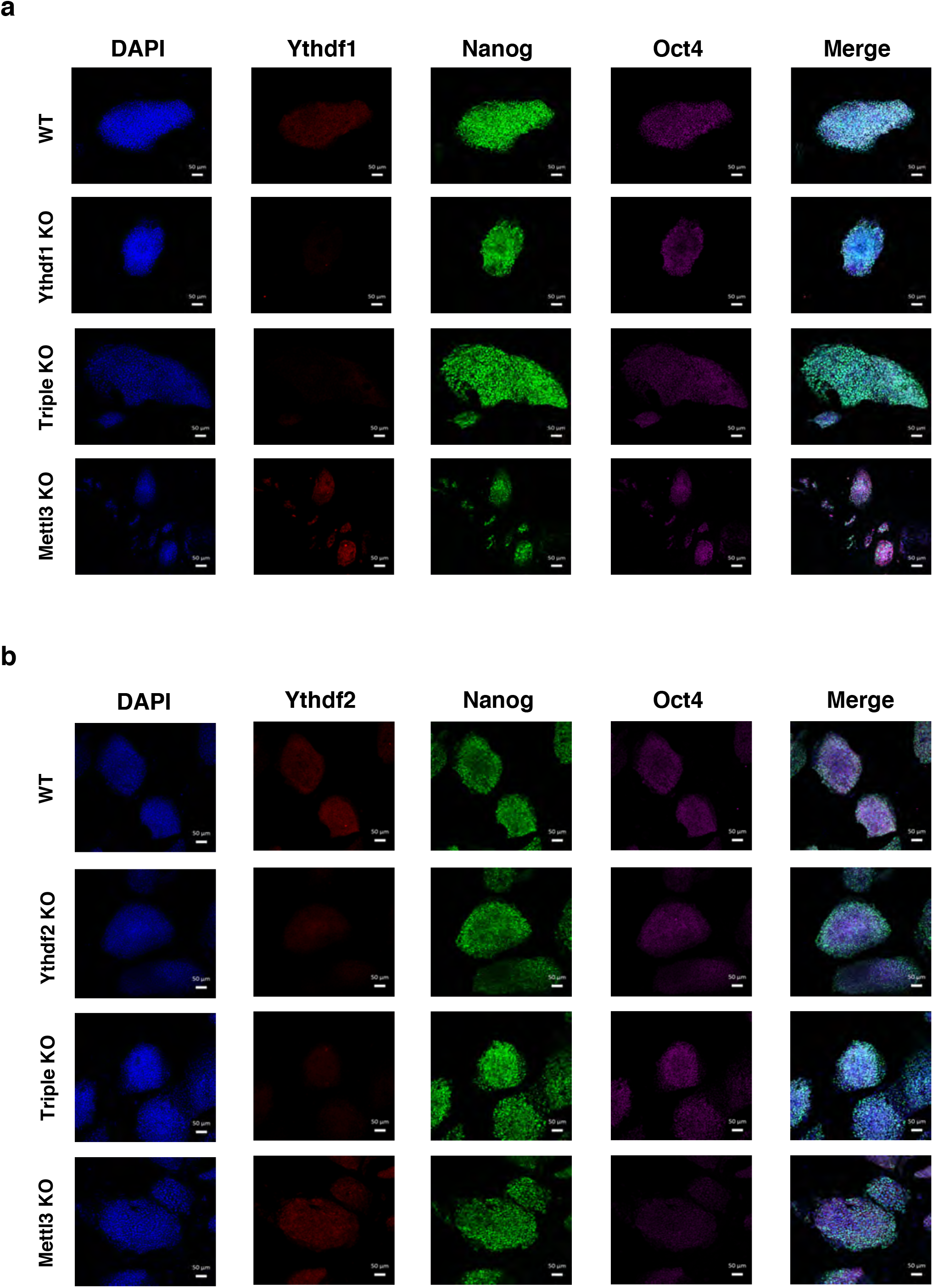
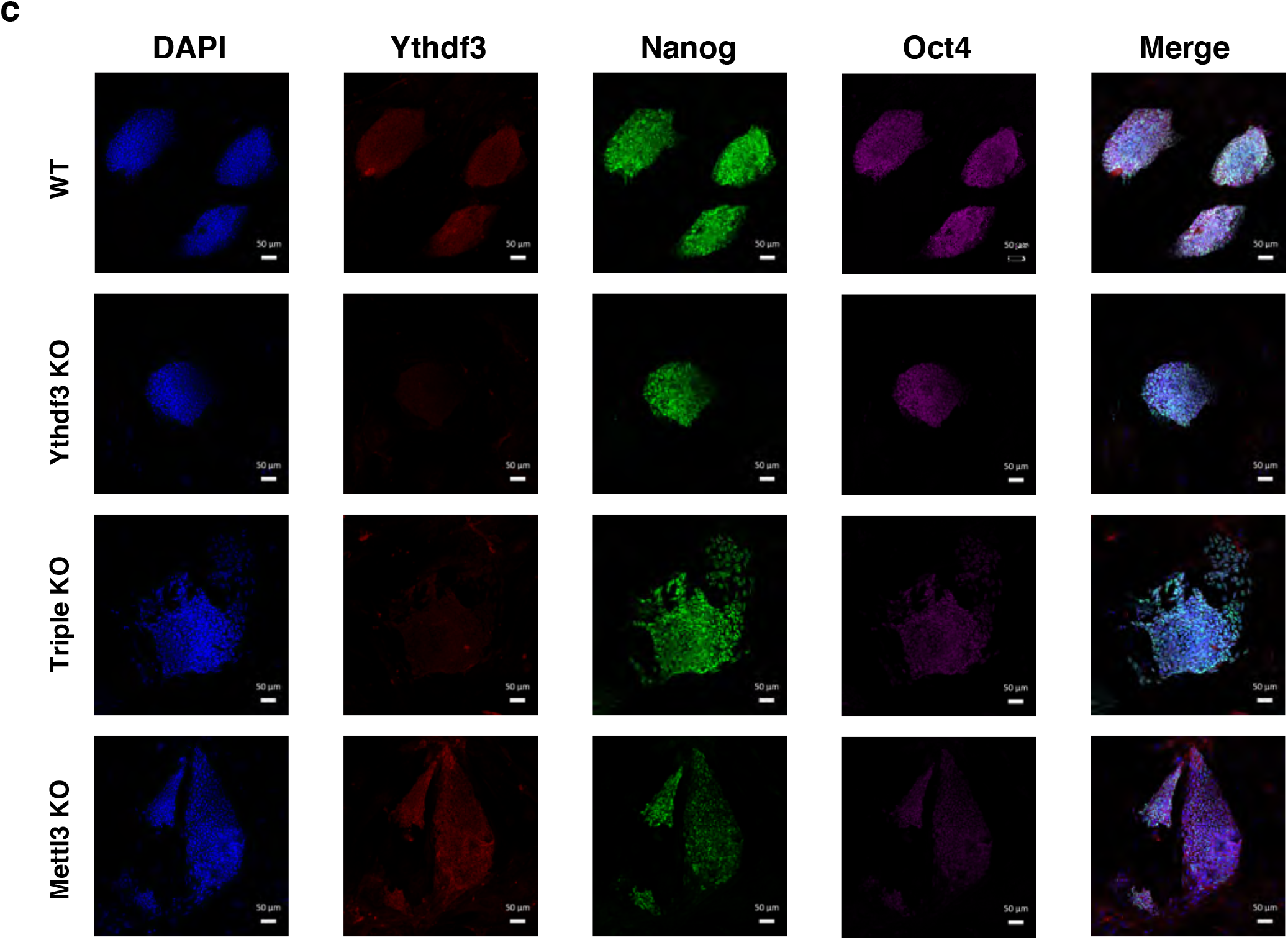
Immunostaining of KO mESC lines. a)□ Immunostaining of Ythdf1 (red), Nanog (green), Oct4 (purple) and DAPI (blue) in WT, Ythdf1-KO, Triple-KO and Mettl3-KO cells. b)□ Immunostaining of Ythdf2 (red), Nanog (green), Oct4 (purple) and DAPI (blue) in WT, Ythdf2-KO, Triple-KO and Mettl3-KO cells. c)□ Immunostaining of Ythdf3 (red), Nanog (green), Oct4 (purple) and DAPI (blue) in WT, Ythdf3-KO, Triple-KO and Mettl3-KO cells.

**Figure S10.**
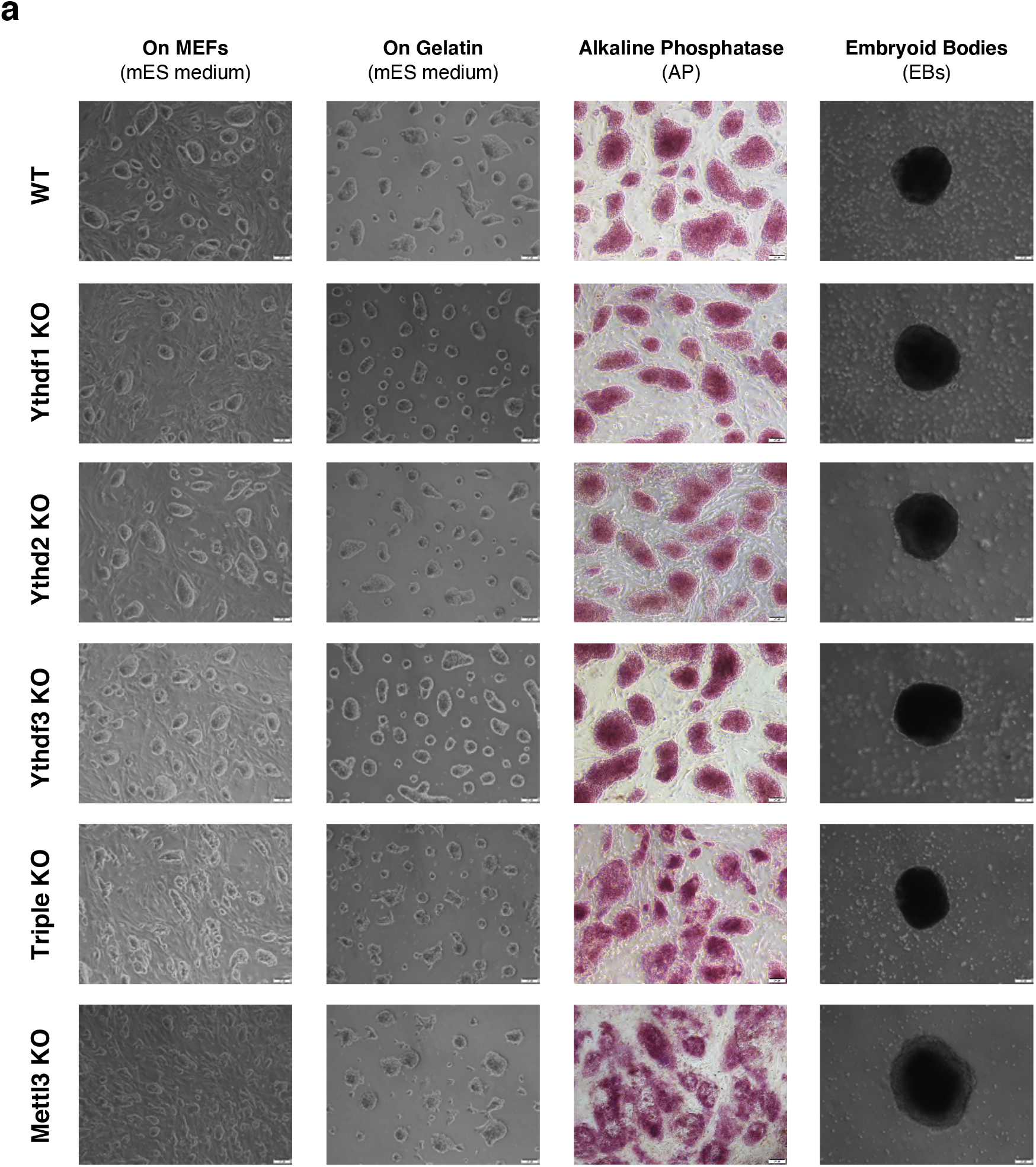
Morphology of knockout mESC lines. a)□ Phase and alkaline phosphatase (AP) staining of WT, single-KOs and Triple-KO mESCs, and phases of their EBs.

**Figure S11.**
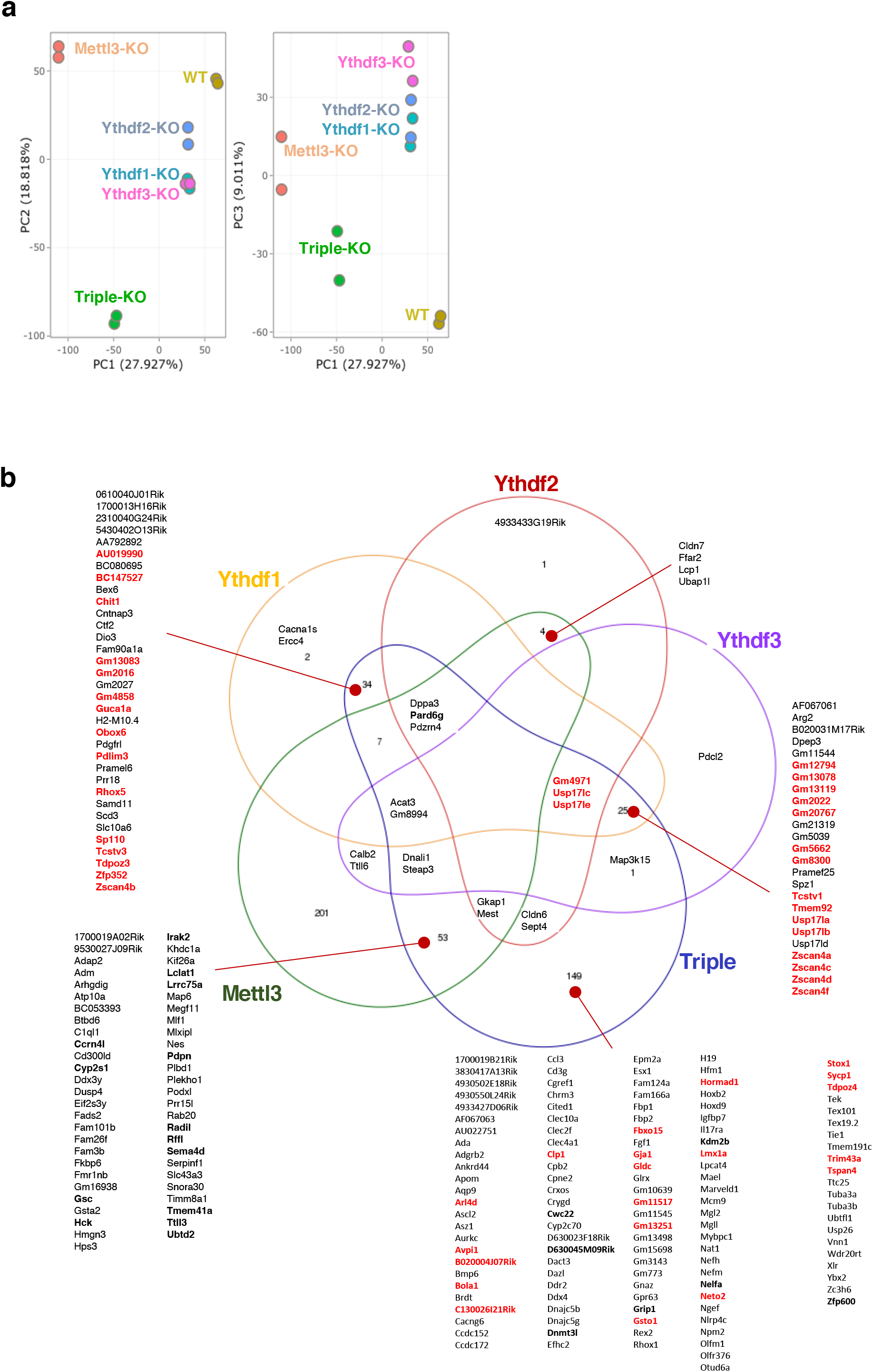
Overlap of ESC signatures. a)□ PCA clustering of KO and WT mESCs samples, showing that in PC1, single reader KO samples are closer to WT, compared to triple-KO and Mettl3-KO. b)□ Overlap between upregulated gene signatures, measured in Ythdf single-KO and triple-KO, and in Mettl3-KO. Genes that are m^6^A-methylated are bold. Genes that are two-cell markers are highlighted in red.

**Figure S12.**
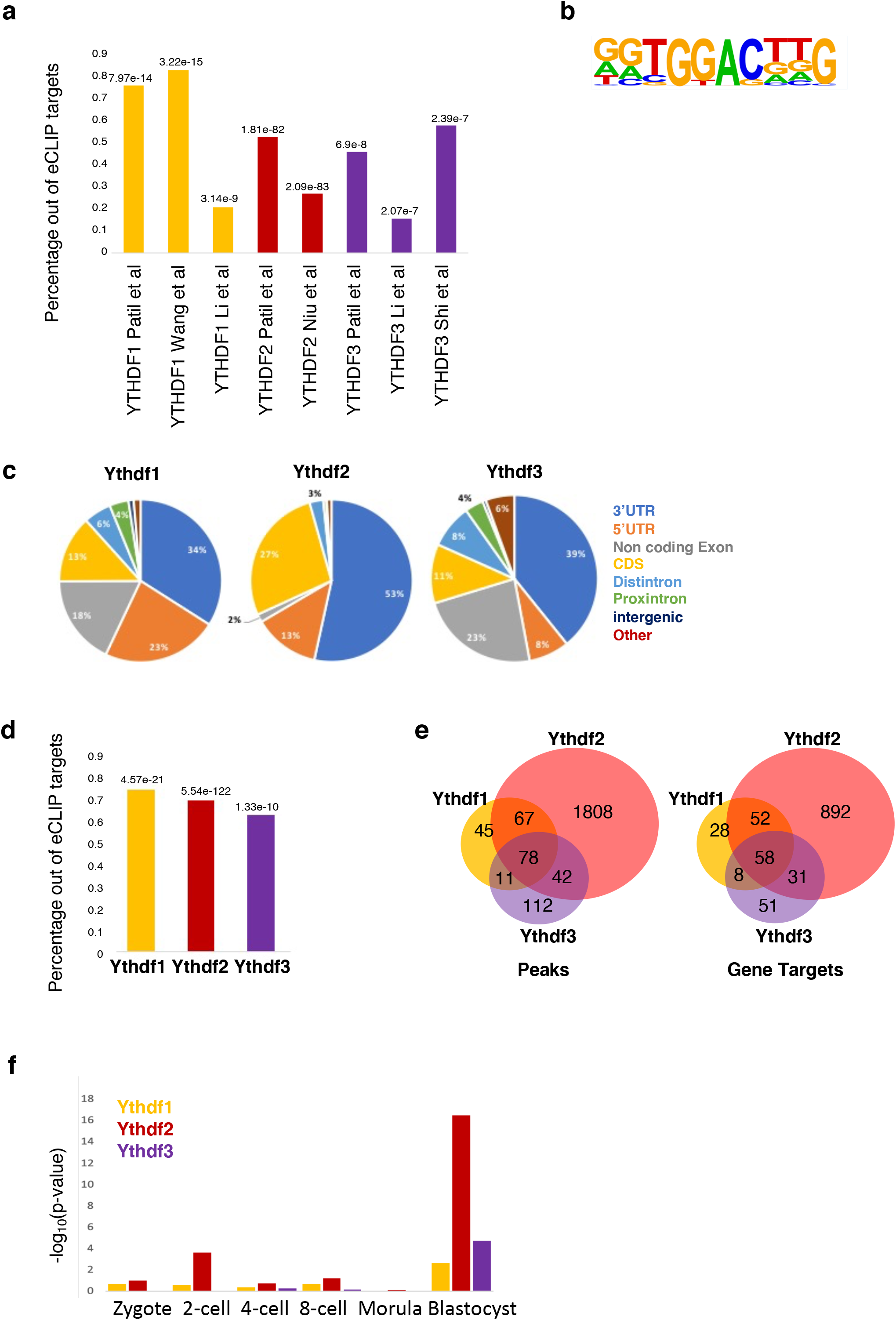
CLIP data evaluation. a)□ Targets of Ythdf1, Ythdf2 and Ythdf3 highly overlap targets that were published before in human cancer cell lines. b)□ Sequence logo of the GGACT containing motif which appears in 14% of YTHDF targets (enrichment fold-change 1.89, p<1e-24). c)□ Distribution of Ythdf peaks in various genomic entities, showing that the three readers have a tendency to bind 3' UTR, particularly Ythdf2. d)□ Significant overlap of Ythdf targets, with m^6^a-methylated genes. e)□ Significant overlap between Ythdf1, Ythdf2 and Ythdf3 targets f)□ Enrichment of Ythdf targets that were identified in mESCs, to early embryo genes (Gao et al. 2017), showing significant overlap with blastocyte genes.

**Figure S13.**
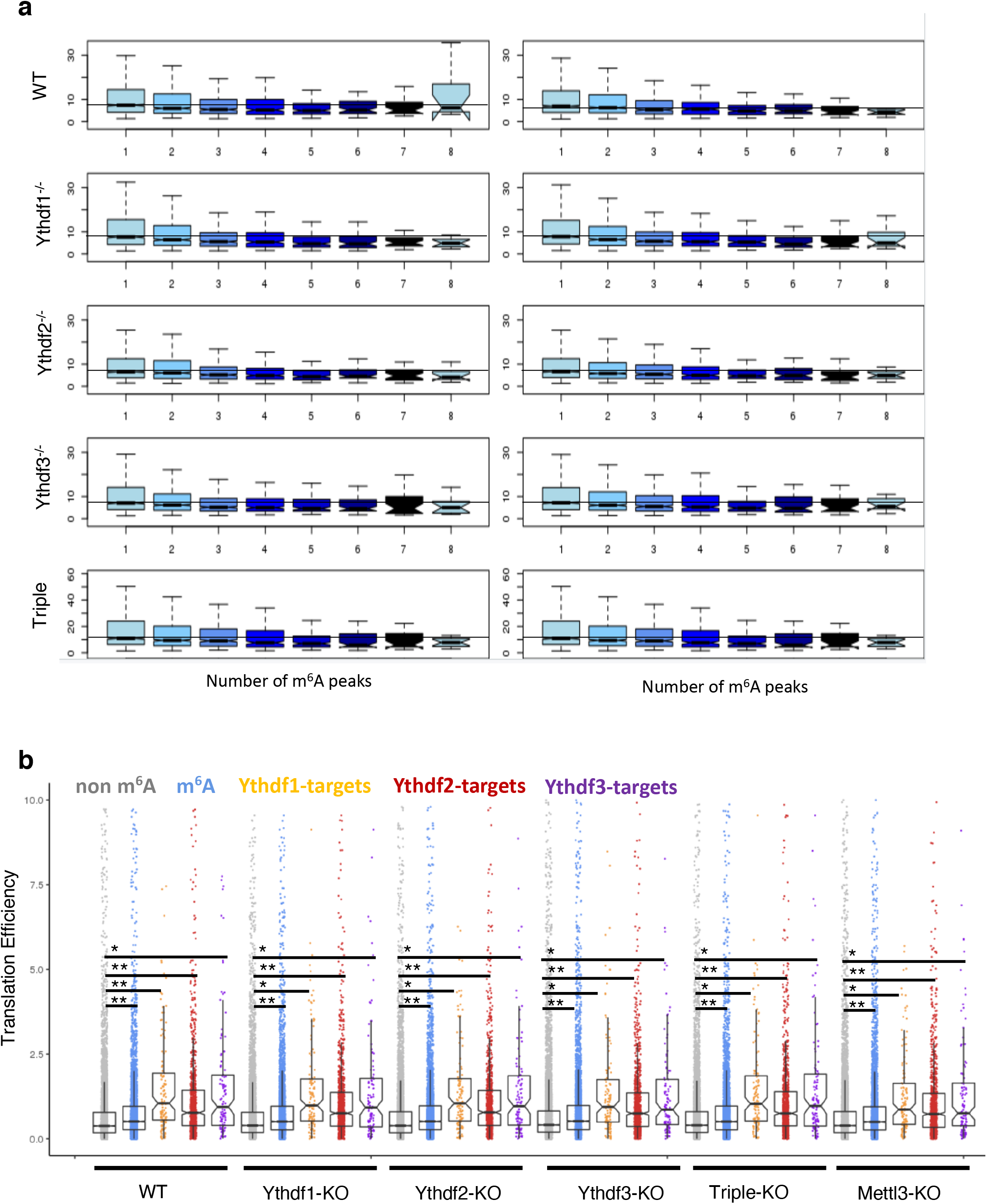
Half-life as a function of number of m^6^A peaks. a)□ The half-life of m^6^A genes is plotted as a function of m6A peak number in the transcript, showing a slight yet significant decrease in half-life (shorter), as the number of m^6^A peaks increase. b)□ Distribution of translation efficiencies of gene groups in the different samples (bottom). Showing that m^6^a-methylated genes and Ythdf targets are translated in a higher efficiency, consistently across samples, compared to non-methylated genes (** p<1e-15, * p<1e-6, Kolmogorov-Smirnov test). Mettl3 and Ythdf KO hardly affect translation efficiency.

## Supplementary Table Legends

**Table S1.** Differentially expressed genes in gametogenesis KO experiments: Mettl3-KO oocytes, and Ythdf2-KO oocytes and spermatoids, compared to matched controls.

**Table S2.** Differentially expressed genes in mESCs, that carry a single-KO (Ythdf1, Ythdf2, Ythdf3 or Mettl3) or triple-KO (Ythdf1/2/3), compared to WT controls.

**Table S3.** eCLIP binding targets of Ythdf1, Ythdf2 and Ythdf3, measured in mESCs.

**Table S4.** Normalized expressed along with transcript half-life, calculated for each gene in the single-KO (Ythdf1, Ythdf2, Ythdf3), triple-KO Ythdf1/2/3, and WT control.

## Materials and Methods

### Stem cell lines and cell culture

Maintenance of WT or Mutant murine ESCs was conducted as described previously **(Geula et al. 2015).** Briefly, mESCs expansion was carried out in 500 mL of High-glucose DMEM (ThermoScientific), 15% USDA certified fetal bovine serum (FBS - Biological Industries), 1 mM L-Glutamine (Biological Industries), 1% nonessential amino acids (Biological Industries), 0.1 mM β-mercaptoethanol (Sigma), 1% penicillin-streptomycin (Biological Industries), 1% Sodium-Pyruvate (Biological Industries), 10μg recombinant human LIF (Peprotech). Cells were maintained in 20% O_2_ conditions on irradiation inactivated mouse embryonic fibroblast (MEF) feeder cells, and were passaged following 0.25% trypsinization. For RNA extraction, cells were grown on Gelatin for three passages in FBS free N2B27-based media **(Gafni et al. 2013)**. Briefly, 500mL of N2B27 media was produced by including: 250 mL DMEM:F12 (ThermoScientific), 250 mL Neurobasal (ThermoScientific), 5 mL N2 supplement (Invitrogen; 17502048 or in-house prepared), 5 mL B27 supplement (Invitrogen; 17504044), 1 mM L-Glutamine (Biological Industries), 1% nonessential amino acids (Biological Industries), 0.1 mM β-mercaptoethanol (Sigma), penicillin-streptomycin (Biological Industries). Naïve conditions for murine ESCs included 10μg recombinant human LIF (Peprotech) and small-molecule inhibitors CHIR99021 (CH, 3 μM-Axon Medchem) and PD0325901 (PD, 1 μM - Axon Medchem) termed 2i.

### Generation of Ythdf1, Ythdf2 and Ythdf3 knock-out murine ESC lines via CRISPR/Cas9

In order to knock out the Ythdf genes in mES cells, oligos for gRNAs were cloned into px335 vector (Addgene#42335 encoding SpCas9 nickase. A pair of unique gRNA sequences for each gene were chosen with the help of the Zhang Lab website http://www.genome-engineering.org/crispr so that to leave 20-30bp offset between the CRISPR target sites. 100 μg of resulting constructs and 10 μg of GFP expressing vector were electroporated into V6.5 mESCs. 3 days later, GFP expressing cells were sorted by FACS and seeded at low density. 9 days after seeding, colonies were picked and their DNA was analyzed by High Resolution Melt assay (HRM) using MeltDoctor reagent (Life Technologies). The clones that showed reduced Tm for the targeted locus were expanded and sequenced to confirm mutations.

gRNA list for knocking out ythdf genes in mES cells:

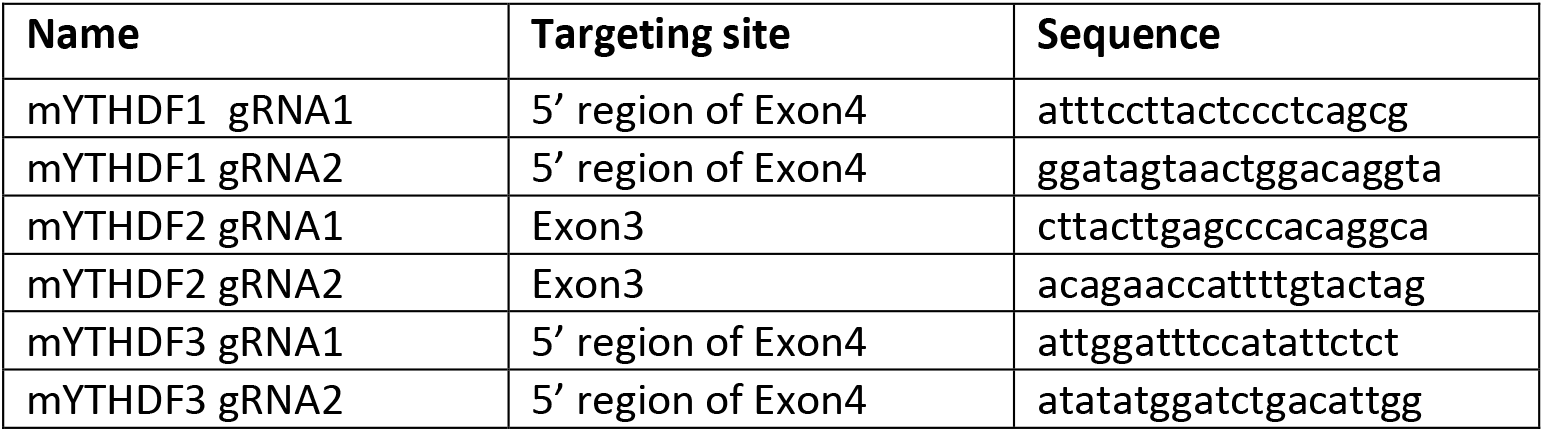

### Generation of Ythdf1, Ythdf2 and Ythdf3 knock-out mouse strains via CRISPR/Cas9

The gRNA sequences were designed with the help of the Zhang Lab website http://www.genome-engineering.org/crispr. For Ythdf1 and Ythdf3 genes single gRNAs were chosen targeting exon3. For Ythdf2 gene, we have designed a pair of gRNAs flanking exon4. The deletion of this exon creates out-of-frame mutation in the coding sequence. Targeting Ythdf genes in mouse single cell embryos was performed as described in **(Yang et al. 2014)**.

Briefly, Cas9 and respective gRNA coding sequences tagged with T7 promoter were transcribed using mMESSAGE mMACHINE T7 ULTRA kit and MEGA shortscript T7 kit, then purified with MEGA clear kit (all the kits were from Thermo Fisher Scientific). CB6F1 (C57BL/6 × BALB/c) and ICR mice strains were used as embryo donors and foster mothers, respectively. Superovulated CB6F1 mice (8-10 weeks old) were mated to CB6F1 stud males, and fertilized embryos were collected from oviducts. Cas9 mRNAs and sgRNA (50 ng/μl) was injected into the cytoplasm of fertilized eggs with well recognized pronuclei in M2 medium (Sigma). The injected zygotes were cultured in KSOM with amino acids (Sigma) at 37°C under 5% CO2 in air until blastocyst stage by 3.5 days. Thereafter, 15-25 blastocysts were transferred into uterus of pseudopregnant ICR females at 2.5 days post coitum (dpc). Mutated animals were screened for deletions by sequencing the targeted loci. Ythdf^+/−^ animal were backcrossed with C57BL/6 mice for 2 generations before mating in order to generate Ythdf^−/−^ knockout mice.

gRNAs for knocking out ythdf genes in mice:

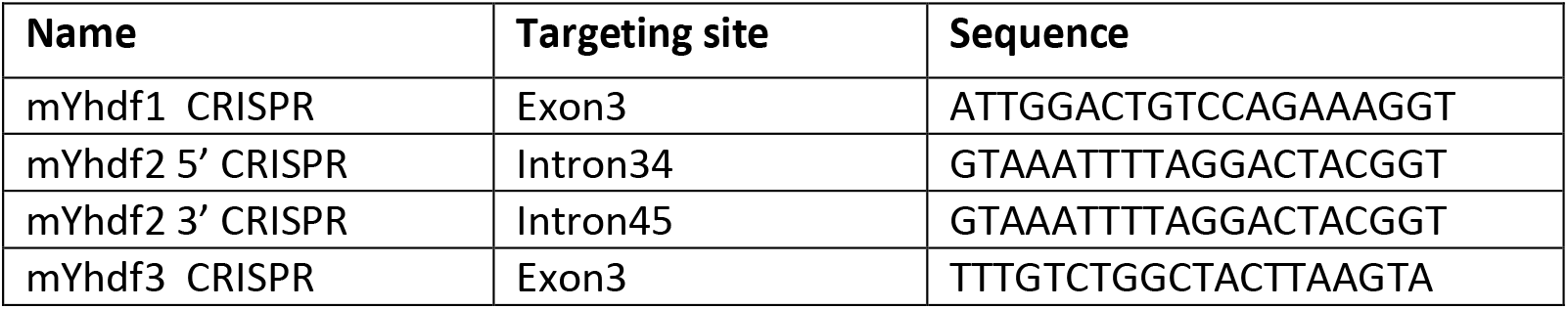

### Generation of Mettl3 conditional-knockout mouse model

Stem cell lines and mice deficient for Mettl3 were generated by targeted disruption of the endogenous Mettl3 locus via homologous recombination. The targeting strategy and construct Knockout Mouse Project repository (Mettl3:tm1a(KOMP)Wtsi) introduced loxP sites spanning the fourth exon that would result in an out-of-frame and truncated product upon deletion and introduced a LacZ reporter cassette driven by the endogenous Mettl3 promoter. 50μg DNA of the targeting construct was linearized and electroporated into V6.5 ESC line that were then subjected to selection with G418 (300microg/ml) After 10 d of selection, resistant clones were analyzed for correct targeting. Mettl3 f/f floxed ESC were injected to BDF1 host blastocyst and chimeric mice were generated. Chimeric male mice were mated with C57BL/6 females. F1 offspring were screened for germline transmission by agouti coat color and validation via PCR of LacZ transgene reporter. In order to remove Neomycin and lacz cassette F1 offspring were mated with Rosa26-FlpE mice (Jackson Laboratory Stock#: 003946) and offspring pups were validated for the removal of LacZ transgene. The mice were crossed to C57BL/6 for 3 generations before used for any experiment.

### Generation of Mettl3^flox/flox^ Cre+ knock-out mice

Cre+ mice were crossed with Mettl3^flox/flox^ mice to generate Mettl3^flox/flox^ Cre+ mice, as detailed in **figure S2b**. The following Cre mice were used: PRM1-Cre (Jax#003328), Stra8-Cre (Jax#017490), Vasa-Cre (Jax#006954) and ZP3-cre+ (Jax#003651).

### Western blot analysis

Cells were harvested, and whole cell protein was extracted by lysis buffer, containing 150 mM NaCl, 150 mM Tris-Hcl (PH = 7.4), 0.5% NP40, 1.5 mM MgCl2, 10% Glycerol. Protein’s concentration was determined by BCA Kit (ThermoScientific). SDS/PAGE was performed according to Laemmli and transferred to nitrocellulose membranes for immunostaining. Membranes containing the transferred proteins were blocked with 5% (w/v) non-fat dried skimmed milk powder in PBST, and then incubated with primary antibody in 5% BSA in PBST (overnight, 4°C). Secondary antibodies used were Peroxide-conjugated AffiniPure goat anti-rabbit (1:10,000, 111-035-003; Jackson ImmunoResearch). Blots were developed using SuperSignal West Pico Chemiluminescent substrate (ThermoScientific, #34080). The following primary antibodies were used: Ythdf2 (AVIVA SYSTEMS BIOLOGY, ARP67917_P050), Ythdf3 (Santa Cruz, SC-87503), Cnot1 (Proteintech, 14276-1-A) and Hsp90 (Epitomics, 1492-1).

### Real Time (RT)-PCR analysis

Total RNA was extracted from the cells using Trizol. 1 μg of RNA was then reverse transcribed using High-Capacity cDNA Reverse Transcription Kit (Applied Biosystems). Quantitative PCR was performed with 10ng of cDNA, in triplicates, on Viia7 platform (Applied Biosystems), using Fast SYBR®Master Mix (Applied Biosystems). Error bars indicate standard deviation of triplicate measurements for each measurement. The primers used for amplification are indicated in the primers table.

Primers list:

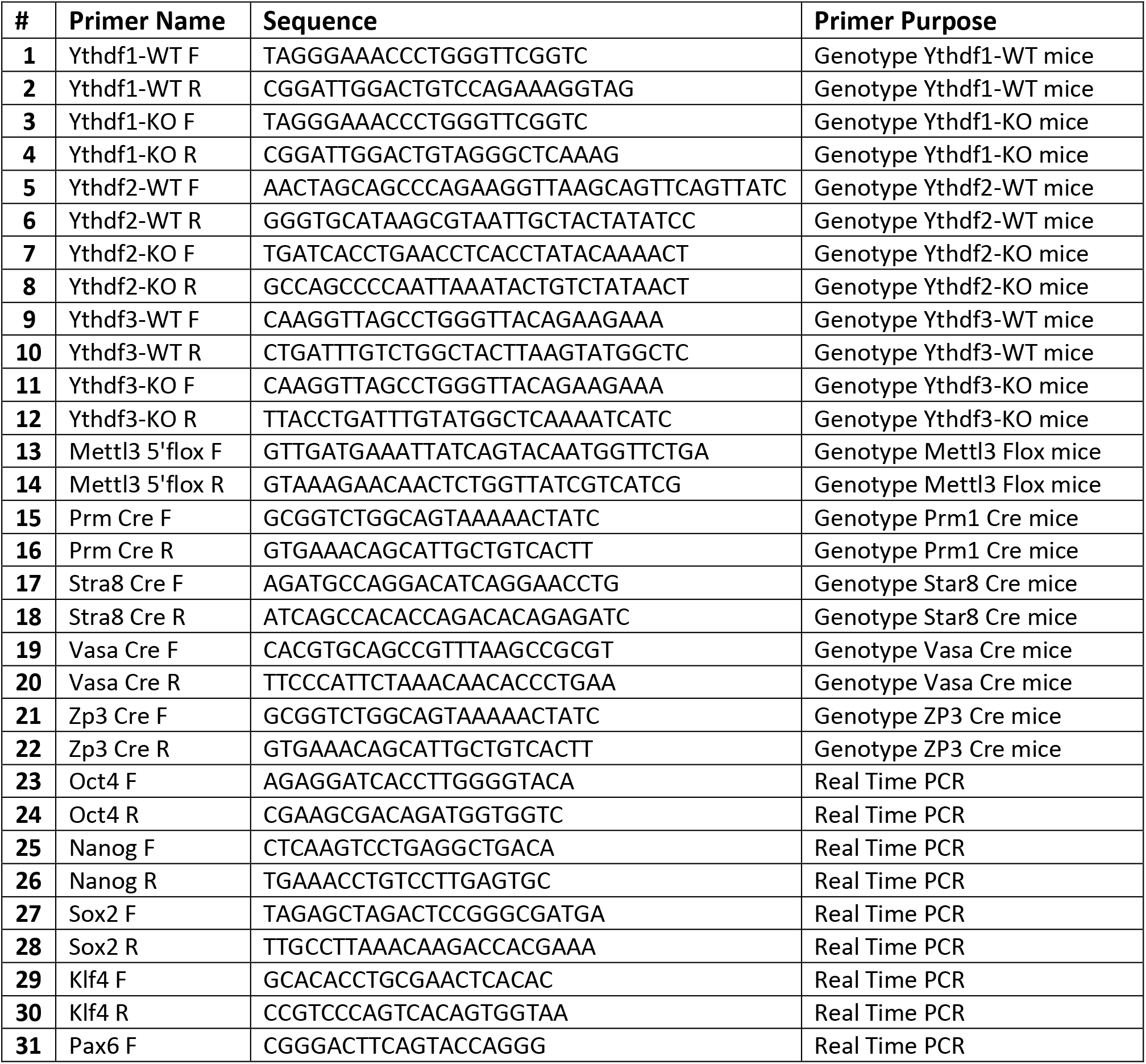
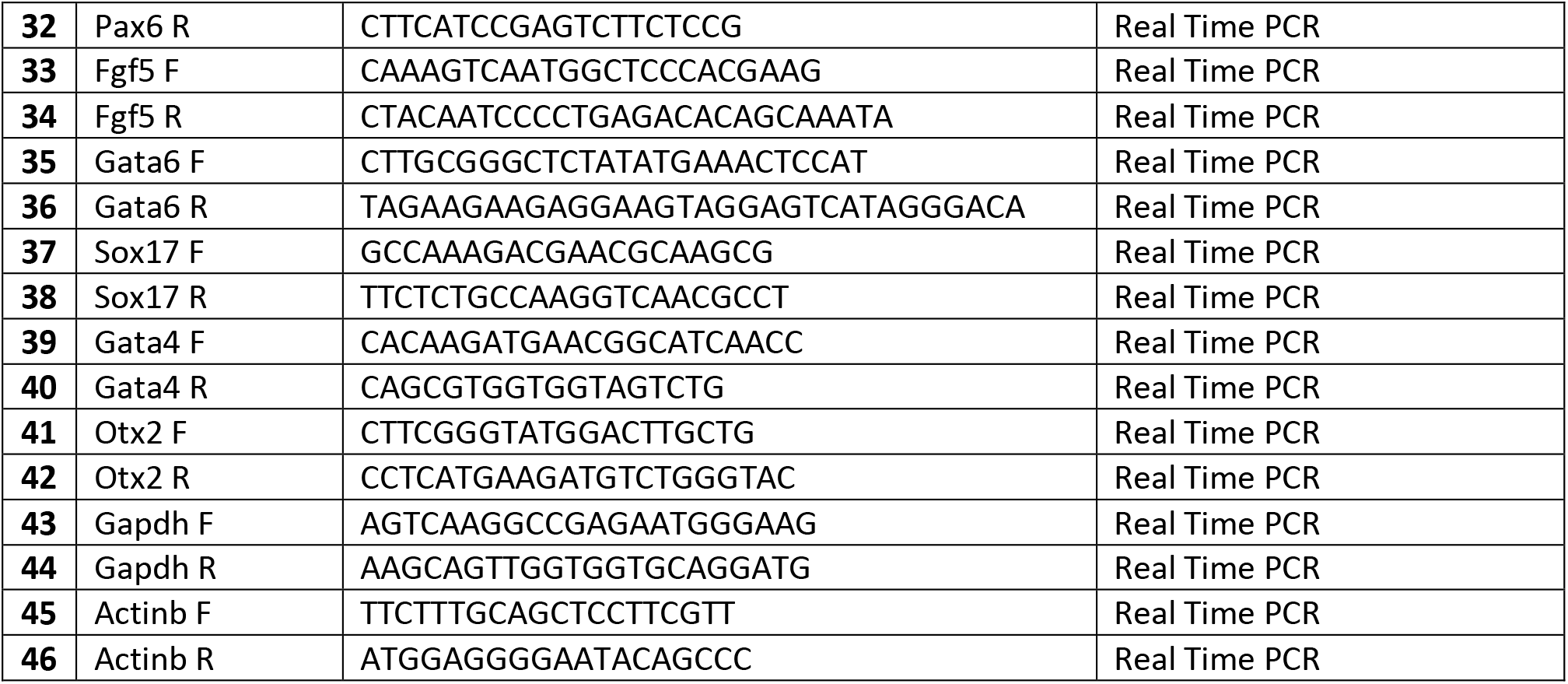

### Embryoid bodies and teratoma formation

For embryoid body (EB) *in vitro* differentiation assay, 5×10^6^ ESCs were disaggregated with trypsin and transferred to non-adherent suspension culture dishes, and cultured in MEF medium (DMEM supplemented with 1% L-Glutamine, 1% Non-essential amino acids, 1% penicillin-streptomycin, 1% Sodium-Pyruvate and 15% FBS, doesn’t contain Lif or 2i) for 8-10 days (time points were always matched with control cells). Media replacement was carried out every 2 days. For teratoma formation, 5×10^6^ ESCs were injected subcutaneously to the flanks of immune deficient NSG mice. After 4-6 weeks, all injected mice were sacrificed and the tumor mass extracted and fixed in 4% para-formaldehyde over-night. Slides were prepared from the paraffin embedded fixed tissues, which were next Hematoxylin & Eosin stained and inspected for representation of all three germ layers.

### Histology

Ovaries and testis were fixed overnight in 4% PFA overnight at 4°C. The fixed tissues were washed with 25%, 50%, and 70% ethanol, embedded in paraffin, and sectioned in 4 μm thickness.

### Oocyte isolation and immunostaining

Female mice (5-8-week old ICR) were injected with 5 i.u. of pregnant mare serum gonadotropin (PMSG) (Sigma), followed by injection of 5 i.u. of human chorionic gonadotrophin (hCG) (Sigma) 46 hours later. Mouse oocytes were extracted from the oviduct by flushing the oviduct with M2 media 24 hours after hCG injection. Somatic cells were removed from the oocytes by gentle pipetting in M2 media supplemented with hyaluronidase (Sigma). Oocytes were transferred to an embryological watch-glass and fixed with 4% PFA EM grade (Electron Microscopy Sciences) in PBS at 4°C over-night. Next, oocytes were washed 3 times in PBS (5 minutes each), permeabilized in PBS with 0.3% Triton X-100 for 30 minutes, blocked with 2% normal donkey serum/0.1% BSA/0.01% Tween-20 in PBS for 1 hour at room temperature (RT), and incubated over-night at 4°C with primary antibodies diluted in blocking solution. Oocytes were rinsed 3 times for 15 minutes each in blocking solution, incubated for 1 hour at room temperature with secondary antibodies diluted 1:500 in blocking solution, counterstained with DAPI (1 μg/ml in PBS) for 5 minutes, and washed with PBS/0.01% Tween-20 for 5 minutes 3 times. Finally, oocytes were mounted in 96 well glass bottom plates for confocal imaging. The following primary antibodies were used: Mouse monoclonal anti-acetylated α-Tubulin (Santa Cruz; sc-23950), Ythdf1 (Proteintech, 17479-1-AP), Ythdf2 (AVIVA SYSTEMS BIOLOGY, ARP67917_P050) and Ythdf3 (Santa Cruz, SC-87503).

### Flushing oocytes

For the collection of oocytes by flushing, female mice were injected with 5 i.u. of pregnant mare serum gonadotropin (PMSG), and after 46 hours with 5 i.u. of human chorionic gonadotropin (hCG). Oocytes were extracted from the oviduct 24 hours after the hCG injection. Next, somatic cells were removed from the oocytes by gentle pipetting in M2 media supplemented with hyaluronidase.

### Poking ovaries

For the collection of GV oocytes, female mice were injected with 5 i.u. of pregnant mare serum gonadotropin (PMSG). After 48 hours, the ovaries were punctured in M2 media, using tweezers. Next, somatic cells were removed from the oocytes by gentle pipetting in M2 media supplemented with hyaluronidase.

### Immunofluorescence staining

Cells subjected to immunofluorescence staining were washed three times with PBS and fixed with 4% paraformaldehyde for 10 minutes at room temperature. Cells were then washed three times with PBS and blocked for 15 min with 5% FBS in PBS containing 0.1% Triton X-100. After incubation with primary antibodies (Over-night, 4°C in 5% FBS in PBS containing 0.1% Tween20) cells were washed three times with PBST (PBS containing 0.1% Tween20) and incubated for one hour at room temperature with fluorophore-labelled appropriate secondary antibodies purchased from Jackson ImmunoResearch. Next, cells were washed and counterstained with DAPI (1 μg/ml, 0215754; MP Biomedical), mounted with Shandon Immu-Mount (ThermoScientific, 9990412) and imaged. All images were collected on LSM700 confocal microscope and processed with Zeiss ZenDesk and Adobe Photoshop CS4 (Adobe Systems, San Jose, CA).

Sections subjected to immunofluorescence staining were rehydrated, treated with antigen retrieval, rinsed in PBS for 5 min, and permeabilized in 0.1% Triton X-100 in PBS, then blocked in Blocking solution (5% normal donkey serum in PBST) in humidified chamber for 1h at RT. Slides were then incubated in the appropriate primary antibody diluted in blocking solution at 4°C overnight. Next, sections were washed three times in PBST, incubated with appropriate fluorochrome-conjugated secondary antibodies diluted in blocking solution for one hour at room temperature in the dark, washed once in PBS, counterstained with DAPI for 10 min, rinsed twice in PBS, mounted with Shandon Immu-Mount (ThermoScientific, 9990412) and imaged. All images were collected on LSM700 confocal microscope and processed with Zeiss ZenDesk and Adobe Photoshop CS4 (Adobe Systems, San Jose, CA). The following primary antibodies were used: Ythdf1 (Proteintech, 17479-1-AP), Ythdf2 (AVIVA SYSTEMS BIOLOGY, ARP67917_P050), Ythdf3 (Santa Cruz, SC-87503), Mettl3 (Proteintech Group 15073-1-AP), Nanog (Bethyl, A300-397A or eBioscience, 14-5761), Esrrb (R&D systems, PP-H6705-00), Oct4 (Santa Cruz, SC9081 or SC5279), Foxa2 (Santa Cruz, sc-6554), Tuj1 (BioLegend, 801202), Tubulin (Santa Cruz, sc-23950). Throughout the manuscript, experimental and control samples were handled for staining, exposure and analysis under identical conditions simultaneously to eliminate variability or bias.

### Alkaline phosphatase (AP) staining

Alkaline phosphatase (AP) staining was performed with AP kit (Millipore SCR004) according to manufacturer protocol. Briefly, cells were fixated using 4% PFA for 2 minutes, and later washed with TBST. The reagents were then added to the wells, followed by an incubation of 10 minutes in RT.

### Tetra complementation (4n) of mouse embryo

4n tetraploid complementation assay was performed by flushing BDF2 embryos at the two-cell stage, and subsequently allowing the embryos to develop until the blastocyst stage. At day 3.5 they were used for PSC micro-injection of triple-KO cell line and its corresponding WT cell line. Embryos were recovered for analysis at E7.5 during development. Embryos were subjected to H&E staining and were observed for developmental defects. All animal studies were conducted according to the guideline and following approval by the Weizmann Institute Institutional Animal Care and Use Committee.

### RNA stability assay

For RNA stability assay 5×10^5^ cells of each cell type were plated on a gelatin-coated 6 cm plate. 48 hours later, the media was replaced with fresh media containing 5 μM Actinomycin-D for the inhibition of mRNA transcription. Cell samples were harvested at the indicated time points (0, 4 and 8 hours) and total RNA was extracted using Trizol, followed by 3’ Poly A-RNA-seq library preparation as previously described (**Geula et al. 2015)**.

### Male Germ cell isolation

Male germ cell populations were isolated using FACS as previously describes **(Bastos et al. 2005; Mahadevaiah et al. 2001; DiGiacomo et al. 2013).** Total RNA was isolated from FACS-sorted round spermatids, from WT control and Ythdf2 KO, using Trizol. The RNA used for RNA-seq (described below)□.

### RNA-seq library preparation

Total RNA was extracted from the indicated mESC cultures using Trizol and treated with DNase to avoid DNA contamination. Polyadenylated RNA was purified using Dynabeads mRNA purification kit (Invitrogen, Cat #61006), followed by library preparation using ScriptSeq v2 RNA-seq Library Preparation Kit (Illumina) according to manufacturer’s instruction.

Male germ cell populations were isolated using FACS as previously described **(Bastos et al. 2005; Mahadevaiah et al. 2001; DiGiacomo et al. 2013)**. Total RNA was extracted using Trizol and purified using rRNA depletion (Ribo-Zero rRNA removal Kit, Illumina), followed by library preparation using ScriptSeq V2 RNA-seq Library Preparation Kit (Illumina) according to manufacturer’s instruction.

### SMART-seq2 library preparation

Library was prepared according to SMART-seq2 protocol as previously described **(Picelli et al. 2014),** with few changes: oocytes from each mouse were collected in 3 ul M2 and added to 7.9 ul lysis buffer. Additional 1.1 ul of DDW was added prior to the library preparation.

### Ribosome profiling & analysis

Ribosome binding profiles in ESCs were measured in WT and KO conditions. For ribosome profiling cells were treated with Cycloheximide as previously described (McGlincy and Ingolia 2017; Ingolia et al. 2009). Cells were then lysed in lysis buffer (20mM Tris 7.5, 150mM NaCl, 15mM MgCl2, 1mM dithiothreitol) supplemented with 0.5% triton, 30 U/ml Turbo DNase (Ambion) and 100μg/ml cycloheximide, ribosome protected fragments were then generated as previously described(McGlincy and Ingolia 2017).

Reads were pre-processed by trimming their linker (sequence CTGTAGGCACCATCAAT) and polyA removal with cutadapt. Reads were aligned to mouse genome version mm10 with Bowtie aligner (parameters -v -m 16 -p 8 --max), where only uniquely aligned reads where used for further analyses. Per gene, for translation calculation, reads were counted in the coding region excluding 15 and 6 nucleotides from the beginning and end of each coding sequence (CDS), respectively **(Ingolia et al. 2009; McGlincy and Ingolia 2017)**. Translation Efficiency was measured for each gene g and each condition i as log2(Ribogi/RNAgi). Normalized translation levels (RPKM) are available alongside the raw data, at NCBI GEO series GSE148039.

### 3’ Poly A-RNA-sequencing Analysis

3’-Poly A-RNA-seq was measured from WT, and KO mESCs. KO cell lines: Ythdf1^−/−^, Ythdf2^−/−^, Ythdf3^−/−^, Ythdf1^−/−^Ythdf2^−/−^Ythdf3^−/−^ and Mettl3^−/−^. In each condition 2 biological replicates were generated, and in each replicate, three time points were measured, 0, 4 and 8 hours after Actinomycin-D induction, with two replicates of time points 0 and 8. In addition, similar 3’-Poly A-RNA-seq dataset from previous paper **(Geula et al. 2015)**, including samples from mESCs and mouse Embryoid bodies (EB), of Mettl3^−/−^ and WT, measured in the same time points, was reanalyzed as described herein.

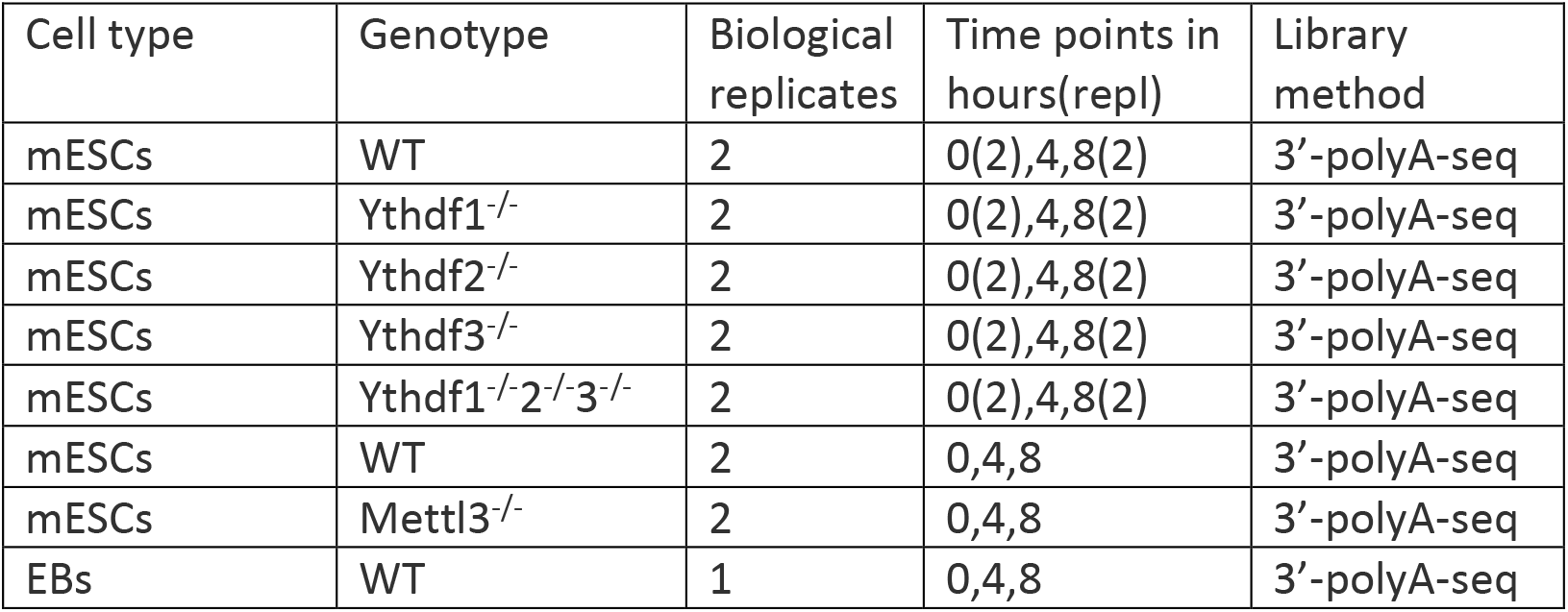
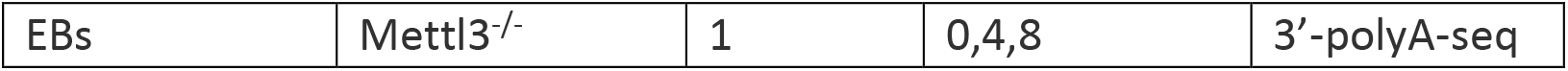

| Cell type | Genotype | Biological replicates | Time points in hours(repl) | Library method |
| --- | --- | --- | --- | --- |
| mESCs | WT | 2 | 0(2),4,8(2) | 3'-polyA-seq |
| mESCs | Ythdf1<sup>-/-</sup> | 2 | 0(2),4,8(2) | 3'-polyA-seq |
| mESCs | Ythdf2<sup>-/-</sup> | 2 | 0(2),4,8(2) | 3'-polyA-seq |
| mESCs | Ythdf3<sup>-/-</sup> | 2 | 0(2),4,8(2) | 3'-polyA-seq |
| mESCs | Ythdf1<sup>-/-</sup>2<sup>-/-</sup>3<sup>-/-</sup> | 2 | 0(2),4,8(2) | 3'-polyA-seq |
| mESCs | WT | 2 | 0,4,8 | 3'-polyA-seq |
| mESCs | Mettl3<sup>-/-</sup> | 2 | 0,4,8 | 3'-polyA-seq |
| EBs | WT | 1 | 0,4,8 | 3'-polyA-seq |
| EBs | Mettl3<sup>-/-</sup> | 1 | 0,4,8 | 3'-polyA-seq |

Reads were aligned to mouse genome version mm10 with Bowtie2 software **(Langmead and Salzberg 2012)** using its default parameters. Gene expression levels were estimated using ESAT software **(Derr et al. 2016)**, and normalized by library size of each sample (FPM, fragments per million reads). To reduce noise, genes were filtered in each sample, to include only genes with at least 3 positive FPM calls (2 in Geula’s dataset), and at least one FPM call > 3 (0.5 in Geula’s dataset), leaving 9K-12K genes in each sample.

3’ polyA RNA-seq values are available alongside the raw data, at NCBI GEO series GSE148039.

### mRNA Half-Life Calculation

The half-life of all genes was calculated according to the following equation: ln(Ci/C0) = -kti, where k is degradation rate, Ci is the mRNA value at time i, and ti is the time interval in hours **(Chen et al. 2008)**. Degradation rate k was estimated for each gene and each sample, from its levels in time points 0h, 4h, 8h (as explained above) using linear regression lm() function in R, on the log transformed levels. Half-life t_1/2_ is ln(2)/k, where k is the degradation rate. Genes which had negative half-life due to slight experimental noise were ignored for the rest of the analysis. Half-life distribution was calculated for each sample, for two groups of genes: non-m^6^A genes, and m^6^A genes which were also bound by either Ythdf1,2 or 3 (m^6^A_YTH). P-value was calculated using Kolmogorov-Smirnov test.

### RNA-Seq analysis

RNA sequencing was measured in mESC, spermatoids and oocytes as detailed in the table below:

| Cell type | Genotype | Biological replicates | Library method | Single/paired-end |
| --- | --- | --- | --- | --- |
| mESCs | WT | 2 | PolyA RNA-seq | Single |
| mESCs | Ythdf1<sup>-/-</sup> | 2 | PolyA RNA-seq | Single |
| mESCs | Ythdf2<sup>-/-</sup> | 2 | PolyA RNA-seq | Single |
| mESCs | Ythdf3<sup>-/-</sup> | 2 | PolyA RNA-seq | Single |
| mESCs | Mettl3<sup>-/-</sup> | 2 | PolyA RNA-seq | Single |
| mESCs | Ythdf1<sup>-/-</sup>2<sup>-/-</sup>3<sup>-/-</sup> | 2 | PolyA RNA-seq | Single |
| GV Oocytes | Mettl3<sup>f/f</sup>Zp3-Cre- | 4 | SMART-seq | paired |
| GV Oocytes | Mettl3<sup>f/f</sup>Zp3-Cre+ | 3 | SMART-seq | paired |
| Post-GV Oocytes | Ythdf2<sup>+/-</sup> | 3 | SMART-seq | paired |
| Post-GV Oocytes | Ythdf2<sup>-/-</sup> | 2 | SMART-seq | paired |
| Round spermatids | Ythdf2<sup>+/+</sup> | 1 | Ribo-zero RNA-seq | paired |
| Round spermatids | Ythdf2<sup>-/-</sup> | 2 | Ribo-zero RNA-seq | paired |

Samples were analyzed using UTAP software **(Kohen et al. 2019)**: Reads were trimmed using cutadapt (Martin 2011) (parameters: -a ADAPTER1 -a “A{10}” -a “T{10}” -A “A{10}” -A “T{10}” – times 2 -q 20 -m 25). Reads were mapped to genome mm10 using STAR **(Dobin et al. 2013)** v2.4.2a (parameters: –alignEndsType EndToEnd, –outFilterMismatchNoverLmax 0.05, –twopassMode Basic). Sample counting was done using STAR, quantifying mm10 RefSeq annotated genes. Further analysis is done for genes having minimum 5 read in at least one sample. Normalization of the counts and differential expression analysis was performed using DESeq2 **(Love et al. 2014)** with the parameters: betaPrior=True, cooksCutoff=FALSE, independentFiltering=FALSE. Raw P values were adjusted for multiple testing using the procedure of Benjamini and Hochberg. Differentially expressed genes were selected with the following parameter: padj <= 0.05, |log2FoldChange| >= 1, baseMean >= 5. PCA and Hierarchical clustering were generated in UTAP software The normalized expression levels are available alongside the raw data, at NCBI GEO series GSE148039.

### Enrichment analysis

Enrichment analysis was done either using GeneAnalytics tool **(Figures S3c, S4f, S6c) (Fuchs et al. 2016)**, or using Fisher exact test **(Figures 5d,S12a,d)**.

### CLIP protocol & CLIP Analysis

Binding targets of Ythdf1, Ythdf2 and Ythdf3 were determined in mESCs using eCLIP method, as described previously (Van Nostrand et al. 2016). 291, 2061 and 306 peaks were identified respectively, mapped to 147, 1034 and 149 genes. Significance of overlap with previous targets and with m^6^A-methylated genes was estimated using Fisher’s exact test **(Figure S12)**. Targets were mapped to human targets in order to test overlap, as previously published targets were measured in human (Patil et al. 2016; Wang et al. 2015; Li et al. 2017; Niu et al. 2013; Shi et al. 2017). Binding Motif was detected using homer software v4.9.1 (http://homer.ucsd.edu/homer/motif/).

## Bibliography

Bailey AS, Batista PJ, Gold RS, Grace Chen Y, de Rooij DG, Chang HY, Fuller MT. 2017. The conserved RNA helicase YTHDC2 regulates the transition from proliferation to differentiation in the germline. Elife 6: E26116.

Dominissini D, Moshitch-Moshkovitz S, Schwartz S, Salmon-Divon M, Ungar L, Osenberg S, Cesarkas K, Jacob-Hirsch J, Amariglio N, Kupiec M, et al. 2012. Topology of the human and mouse m6A RNA methylomes revealed by m6A-seq. Nature 485: 201–206. http://www.ncbi.nlm.nih.gov/pubmed/22575960 (Accessed September 24, 2019).

Du H, Zhao Y, He J, Zhang Y, Xi H, Liu M, Ma J, Wu L. 2016. YTHDF2 destabilizes m 6 A-containing RNA through direct recruitment of the CCR4-NOT deadenylase complex. Nat Commun 7: 12626.

Gao LL, Zhou CX, Zhang XL, Liu P, Jin Z, Zhu GY, Ma Y, Li J, Yang ZX, Zhang D. 2017a. ZP3 is Required for Germinal Vesicle Breakdown in Mouse Oocyte Meiosis. Sci Rep. 7: 41272.

Gao Y, Liu X, Tang B, Li C, Kou Z, Li L, Liu W, Wu Y, Kou X, Li J, et al. 2017b. Protein Expression Landscape of Mouse Embryos during Pre-implantation Development. Cell Rep 21: 3957–3969.

Garcia-Campos MA, Edelheit S, Toth U, Safra M, Shachar R, Viukov S, Winkler R, Nir R, Lasman L, Brandis A, et al. 2019. Deciphering the “m6A Code” via Antibody-Independent Quantitative Profiling. Cell 178: 731–747.

Geula S, Moshitch-Moshkovitz S, Dominissini D, Mansour AA, Kol N, Salmon-Divon M, Hershkovitz V, Peer E, Mor N, Manor YS, et al. 2015. m^6^A mRNA methylation facilitates resolution of naïve pluripotency toward differentiation. Science (80-) 347.

Hartmann AM, Nayler O, Schwaiger FW, Obermeier A, Stamm S. 1999. The interaction and colocalization of Sam68 with the splicing-associated factor YT521-B in nuclear dots is regulated by the Src family kinase p59(fyn). Mol Biol Cell 10: 3909–26.

Heck AM, Wilusz CJ. 2019. Small changes, big implications: The impact of m6A RNA methylation on gene expression in pluripotency and development. Biochim Biophys Acta - Gene Regul Mech 1862: 194402.

Hsu PJ, Zhu Y, Ma H, Guo Y, Shi X, Liu Y, Qi M, Lu Z, Shi H, Wang J, et al. 2017. Ythdc2 is an N6 - methyladenosine binding protein that regulates mammalian spermatogenesis. Cell Res 27: 1115–1127.

Huang T, Liu Z, Zheng Y, Feng T, Gao Q, Zeng W. 2020. YTHDF2 promotes spermagonial adhesion through modulating MMPs decay via m 6 A / mRNA pathway. Cell Death Dis 11: 37.

Ivanova I, Much C, Di Giacomo M, Azzi C, Morgan M, Moreira PN, Monahan J, Carrieri C, Enright AJ, O’Carroll D. 2017. The RNA m6A Reader YTHDF2 Is Essential for the Post-transcriptional Regulation of the Maternal Transcriptome and Oocyte Competence. Mol Cell 67: 1059–1067.

Jain D, Puno MR, Meydan C, Lailler N, Mason CE, Lima CD, Anderson K V., Keeney S. 2018. Ketu mutant mice uncover an essential meiotic function for the ancient RNA helicase YTHDC2. Elife 7: E30919.

Jia G, Fu Y, Zhao X, Dai Q, Zheng G, Yang Y, Yi C, Lindahl T, Pan T, Yang YG, et al. 2011. N6-Methyladenosine in nuclear RNA is a major substrate of the obesity-associated FTO. Nat Chem Biol 7: 885–7.

Kan L, Grozhik A V., Vedanayagam J, Patil DP, Pang N, Lim KS, Huang YC, Joseph B, Lin CJ, Despic V, et al. 2017. The m 6 A pathway facilitates sex determination in Drosophila. Nat Commun 8: 15737.

Kasowitz SD, Ma J, Anderson SJ, Leu NA, Xu Y, Gregory BD, Schultz RM, Wang PJ. 2018. Nuclear m6A reader YTHDC1 regulates alternative polyadenylation and splicing during mouse oocyte development. PLoS Genet 14: e1007412.

Lasman, L, Hanna, JH, Novershtern N. 2020. Role of m6A in Embryonic Stem Cell Differentiation and in Gametogenesis. Epigenomes 4.

Lee H, Bao S, Qian Y, Geula S, Leslie J, Zhang C, Hanna JH, Ding L. 2019. Stage-specific requirement for Mettl3-dependent m6A mRNA methylation during haematopoietic stem cell differentiation. Nat Cell Biol 21: 700–709.

Li A, Chen YS, Ping XL, Yang X, Xiao W, Yang Y, Sun HY, Zhu Q, Baidya P, Wang X, et al. 2017. Cytoplasmic m 6 A reader YTHDF3 promotes mRNA translation. Cell Res.

Li M, Zhao X, Wang W, Shi H, Pan Q, Lu Z, Perez SP, Suganthan R, He C, Bjørås M, et al. 2018. Ythdf2-mediated m6A mRNA clearance modulates neural development in mice. Genome Biol 19: 69.

Lin Z, Hsu PJ, Xing X, Fang J, Lu Z, Zou Q, Zhang KJ, Zhang X, Zhou Y, Zhang T, et al. 2017. Mettl3-/Mettl14-mediated mRNA N 6-methyladenosine modulates murine spermatogenesis. Cell Res 27: 1216–1230.

Liu J, Yue Y, Han D, Wang X, Fu Y, Zhang L, Jia G, Yu M, Lu Z, Deng X, et al. 2014. A METTL3-METTL14 complex mediates mammalian nuclear RNA N6-adenosine methylation. Nat Chem Biol 10: 93–5.

Meyer KD, Saletore Y, Zumbo P, Elemento O, Mason CE, Jaffrey SR. 2012. Comprehensive analysis of mRNA methylation reveals enrichment in 3′ UTRs and near stop codons. Cell 149: 1635–46.

Niu Y, Zhao X, Wu YS, Li MM, Wang XJ, Yang YG. 2013. N6-methyl-adenosine (m6A) in RNA: An Old Modification with A Novel Epigenetic Function. Genomics, Proteomics Bioinforma 11: 8–17.

O’Donnell L, O’Bryan MK. 2014. Microtubules and spermatogenesis. Semin Cell Dev Biol 30: 45–54.

Patil DP, Chen CK, Pickering BF, Chow A, Jackson C, Guttman M, Jaffrey SR. 2016. M6 A RNA methylation promotes XIST-mediated transcriptional repression. Nature 537: 369–373.

Patil DP, Pickering BF, Jaffrey SR. 2018. Reading m6A in the Transcriptome: m6A-Binding Proteins. Trends Cell Biol 28: 113–127.

Pervaiz N, Shakeel N, Qasim A, Zehra R, Anwar S, Rana N, Xue Y, Zhang Z, Bao Y, Abbasi AA. 2019. Evolutionary history of the human multigene families reveals widespread gene duplications throughout the history of animals. BMC Evol Biol 19: 128.

Schwartz S, Agarwala SD, Mumbach MR, Jovanovic M, Mertins P, Shishkin A, Tabach Y, Mikkelsen TS, Satija R, Ruvkun G, et al. 2013. High-Resolution mapping reveals a conserved, widespread, dynamic mRNA methylation program in yeast meiosis. Cell 155: 1409–21.

Shi H, Wang X, Lu Z, Zhao BS, Ma H, Hsu PJ, Liu C, He C. 2017. YTHDF3 facilitates translation and decay of N 6-methyladenosine-modified RNA. Cell Res 27: 315–328.

Shi H, Zhang X, Weng YL, Lu Z, Liu Y, Lu Z, Li J, Hao P, Zhang Y, Zhang F, et al. 2018. m6A facilitates hippocampus-dependent learning and memory through YTHDF1. Nature 563: 249–253.

Stern-Ginossar N, Weisburd B, Michalski A, Le VTK, Hein MY, Huang SX, Ma M, Shen B, Qian SB, Hengel H, et al. 2012. Decoding human cytomegalovirus. Science (80-) 338: 1088–93.

Storm MP, Kumpfmueller B, Thompson B, Kolde R, Vilo J, Hummel O, Schulz H, Welham MJ. 2009. Characterization of the phosphoinositide 3-kinase-dependent transcriptome in murine embryonic stem cells: Identification of novel regulators of pluripotency. Stem Cells 27: 764–75.

Tang C, Klukovich R, Peng H, Wang Z, Yu T, Zhang Y, Zheng H, Klungland A, Yan W. 2017. ALKBH5-dependent m6A demethylation controls splicing and stability of long 3’-UTR mRNAs in male germ cells. Proc Natl Acad Sci U S A 115: E325–E333.

Van Nostrand EL, Pratt GA, Shishkin AA, Gelboin-Burkhart C, Fang MY, Sundararaman B, Blue SM, Nguyen TB, Surka C, Elkins K, et al. 2016. Robust transcriptome-wide discovery of RNA-binding protein binding sites with enhanced CLIP (eCLIP). Nat Methods 13: 508–14.

Wang X, Lu Z, Gomez A, Hon GC, Yue Y, Han D, Fu Y, Parisien M, Dai Q, Jia G, et al. 2014. N 6-methyladenosine-dependent regulation of messenger RNA stability. Nature 505: 117–20.

Wang X, Zhao BS, Roundtree IA, Lu Z, Han D, Ma H, Weng X, Chen K, Shi H, He C. 2015. N6-methyladenosine modulates messenger RNA translation efficiency. Cell 161: 1388–99.

Wang Y, Li Y, Yue M, Wang J, Kumar S, Wechsler-Reya RJ, Zhang Z, Ogawa Y, Kellis M, Duester G, et al. 2018. N 6-methyladenosine RNA modification regulates embryonic neural stem cell self-renewal through histone modifications. Nat Neurosci 21: 195–206.

Wei J, Liu F, Lu Z, Fei Q, Ai Y, He PC, Shi H, Cui X, Su R, Klungland A, et al. 2018. Differential m 6 A, m 6 A m, and m 1 A Demethylation Mediated by FTO in the Cell Nucleus and Cytoplasm. Mol Cell 71: 973–985.

Wojtas MN, Pandey RR, Mendel M, Homolka D, Sachidanandam R, Pillai RS. 2017. Regulation of m 6 A Transcripts by the 3′→5′ RNA Helicase YTHDC2 Is Essential for a Successful Meiotic Program in the Mammalian Germline. Mol Cell.

Xia H, Zhong C, Wu X, Chen J, Tao B, Xia X, Shi M, Zhu Z, Trudeau VL, Hu W. 2018. Mettl3 mutation disrupts gamete maturation and reduces fertility in zebrafish. Genetics.

Xu K, Yang Y, Feng GH, Sun BF, Chen JQ, Li YF, Chen YS, Zhang XX, Wang CX, Jiang LY, et al. 2017. Mettl3-mediated m 6 A regulates spermatogonial differentiation and meiosis initiation. Cell Res 27: 1100–1114.

Zheng G, Dahl JA, Niu Y, Fedorcsak P, Huang CM, Li CJ, Vågbø CB, Shi Y, Wang WL, Song SH, et al. 2013. ALKBH5 Is a Mammalian RNA Demethylase that Impacts RNA Metabolism and Mouse Fertility. Mol Cell 49: 18–29.

## Bibliography

Bastos H, Lassalle B, Chicheportiche A, Riou L, Testart J, Allemand I, Fouchet P. 2005. Flow cytometric characterization of viable meiotic and postmeiotic cells by Hoechst 33342 in mouse spermatogenesis. Cytom Part A 65: 40–9.

Chen CYA, Ezzeddine N, Shyu A Bin. 2008. Chapter 17 Messenger RNA Half-Life Measurements in Mammalian Cells. In Methods in Enzymology, pp. 335–57.

Derr A, Yang C, Zilionis R, Sergushichev A, Blodgett DM, Redick S, Bortell R, Luban J, Harlan DM, Kadener S, et al. 2016. End Sequence Analysis Toolkit (ESAT) expands the extractable information from single-cell RNA-seq data. Genome Res 26: 1397–1410.

DiGiacomo M, Comazzetto S, Saini H, DeFazio S, Carrieri C, Morgan M, Vasiliauskaite L, Benes V, Enright AJ, O’Carroll D. 2013. Multiple Epigenetic Mechanisms and the piRNA Pathway Enforce LINE1 Silencing during Adult Spermatogenesis. Mol Cell 50: 601–8.

Dobin A, Davis CA, Schlesinger F, Drenkow J, Zaleski C, Jha S, Batut P, Chaisson M, Gingeras TR. 2013. STAR: Ultrafast universal RNA-seq aligner. Bioinformatics 29: 15–21.

Fuchs SBA, Lieder I, Stelzer G, Mazor Y, Buzhor E, Kaplan S, Bogoch Y, Plaschkes I, Shitrit A, Rappaport N, et al. 2016. GeneAnalytics: An Integrative Gene Set Analysis Tool for Next Generation Sequencing, RNAseq and Microarray Data. Omi A J Integr Biol 20: 139–51.

Gafni O, Weinberger L, Mansour AA, Manor YS, Chomsky E, Ben-Yosef D, Kalma Y, Viukov S, Maza I, Zviran A, et al. 2013. Derivation of novel human ground state naive pluripotent stem cells. Nature 504.

Ingolia NT, Ghaemmaghami S, Newman JRS, Weissman JS. 2009. Genome-wide analysis in vivo of translation with nucleotide resolution using ribosome profiling. Science (80-) 324: 218–23.

Kohen R, Barlev J, Hornung G, Stelzer G, Feldmesser E, Kogan K, Safran M, Leshkowitz D. 2019. UTAP: User-friendly Transcriptome Analysis Pipeline. BMC Bioinformatics 20: 154.

Langmead B, Salzberg SL. 2012. Fast gapped-read alignment with Bowtie 2. Nat Methods 9: 357–9.

Love MI, Huber W, Anders S. 2014. Moderated estimation of fold change and dispersion for RNA-seq data with DESeq2. Genome Biol 15: 550.

Mahadevaiah SK, Turner JMA, Baudat F, Rogakou EP, De Boer P, Blanco-Rodríguez J, Jasin M, Keeney S, Bonner WM, Burgoyne PS. 2001. Recombinational DNA double-strand breaks in mice precede synapsis. Nat Genet 27: 271–6.

Martin M. 2011. Cutadapt removes adapter sequences from high-throughput sequencing reads. EMBnet.journal.

McGlincy NJ, Ingolia NT. 2017. Transcriptome-wide measurement of translation by ribosome profiling. Methods 126: 112–129.

Picelli S, Faridani OR, Björklund ÅK, Winberg G, Sagasser S, Sandberg R. 2014. Full-length RNA-seq from single cells using Smart-seq2. Nat Protoc 9: 171–81.

Yang H, Wang H, Jaenisch R. 2014. Generating genetically modified mice using CRISPR/Cas-mediated genome engineering. Nat Protoc 9: 1956–68.

